# Deep Learning for Flexible and Site-Specific Protein Docking and Design

**DOI:** 10.1101/2023.04.01.535079

**Authors:** Matt McPartlon, Jinbo Xu

**Affiliations:** The University of Chicago, Department of Computer Science; MoleculeMind Inc. Beijing, China; Toyota Technical Institute at Chicago

## Abstract

Protein complexes are vital to many biological processes and their understanding can lead to the development of new drugs and therapies. Although the structure of individual protein chains can now be predicted with high accuracy, determining the three-dimensional structure of a complex remains a challenge. Protein docking, the task of computationally determining the structure of a protein complex given the unbound structures of its components (and optionally binding site information), provides a way to predict protein complex structure. Traditional docking methods rely on empirical scoring functions and rigid body simulations to predict the binding poses of two or more proteins. However, they often make unrealistic assumptions about input structures, and are not effective at accommodating conformational flexibility or binding site information. In this work, we present DockGPT (Generative Protein Transformer for Docking), an end-to-end deep learning method for flexible and site-specific protein docking that allows conformational flexibility and can effectively make use of binding site information. Tested on multiple benchmarks with unbound and predicted monomer structures as input, we significantly outperform existing methods in both accuracy and running time. Our performance is especially pronounced for antibody-antigen complexes, where we predict binding poses with high accuracy even in the absence of binding site information. Finally, we highlight our method’s generality by extending it to simultaneously dock and co-design the sequence and structure of antibody complementarity determining regions targeting a specified epitope.

## 1 Introduction

The bound configuration of two or more proteins helps regulate many biological processes including signal transduction [1, 2], membrane transport [3, 4], and cell metabolism [5, 6]. The process by which unbound protein chains bind together to form a complex is often controlled by more general protein-protein interactions (PPIs) [7–9], and accordingly, aberrant PPIs are associated with various diseases, including cancer, infectious diseases, and neurodegenerative diseases [10]. The role of PPIs in protein complex formation makes selective targeting of PPIs an essential strategy for drug design and already forms the basis for several established cancer immunotherapies such as monoclonal antibodies [11, 12]. Although most proteins interact with partners to form a complex, experimental methods for determining the structures are often expensive and technically difficult to administer [13, 14]. As a result, protein complexes account for only a small fraction of entries in the Protein Data Bank (PDB) [15], highlighting the need for effective *in silico* methods.

Although it is possible to infer protein complex struc-ture from primary sequence information alone, in many cases, the three-dimensional structures of constituent (unbound) chains have already been experimentally determined. Moreover, extra information such as target binding sites or inter-chain contacts, is readily available in many applications, or can be derived through experimental methods such as cross-linking mass spectrometry [16]. In these scenarios, protein docking methods can be used to predict a complex structure. Despite having many practical applications [17–19], the efficacy of *in silico* protein docking or design methods is ultimately hindered by unrealistic assumptions about input structures, and failure to effectively utilize PPI information such as binding sites and inter-protein contacts.

Current computational methods for protein docking and design typically impose backbone and side-chain rigidity constraints and are trained to utilize specific side-chain interactions or protein backbone placements derived from native complexes which are already optimal for binding [20, 21]. Training computational models on only bound structures – in which binding interfaces match perfectly– is in a sense “starting with the answer.” In the real world, unbound chains typically lack shape complementarity because proteins tend to deform substantially upon binding [22, 23], even for small-molecule ligands [24]. Accounting for backbone and side-chain flexibility can significantly increase the number of sequences that fold to the structure while maintaining the general fold of the protein [25], and is especially important for protein design because mutations in sequence often result in small changes to the backbone structure, [26], and potentially large changes to surrounding side-chain conformations [27].

In addition to overlooking conformational flexibility, current methods tend to either ignore or ineffectively incorporate PPI information. For many applications, it is important to consider interactions as a particular binding site, such as targeting catalytic sites of enzymes, or designing therapeutics to block a specific protein-protein interaction. A salient example is the design of neutralizing antibodies targeting the SARS-CoV-2 S protein which initiates infection upon binding to the human angiotensin-converting enzyme 2 (ACE2) receptor [28, 29]. In most cases, PPIs such as binding sites or interchain contacts are utilized only as a post-processing step, to re-rank or filter out incompatible predictions.

Flexible docking and design of protein complexes presents several challenges for machine learning. First, the 3D geometry of multiple proteins is inherently difficult to represent. The difficulty arises from the fact that spatial relationships between receptor and ligand structures are ambiguous at the input level, yet inter-protein interactions must still be modeled jointly by the learning algorithm. Although several geometric deep-learning approaches offer a way to directly model 3D point clouds, so far only one end-to-end machine learning method has been proposed for general protein docking [30]. This method does not take into consideration backbone flexibility or bindings site information, and suffers from excessive steric clashes in its predictions. Finally, sufficient training data is also scarce. Currently, there is no large dataset consisting of both protein complexes and their unbound components.

In this work, we introduce DockGPT, an end-to-end deep-learning approach to site-specific flexible docking and design. In developing DockGPT, we hypothesized that neural networks could accurately recover protein 3D-coordinates from coarse or incomplete descriptions of their geometry. After affirming this capability, we approached flexible docking in a manner analogous to matrix completion followed by multidimensional scaling. In the matrix completion step, missing entries loosely correspond to inter-chain quantities such as distance and orientation. The imputed representation is then converted to 3D geometry in order to recover the bound complex. This framing allows us to naturally incorporate PPI information as input, in the form of residue-level binding interfaces or interfacial contacts. In addition, removing some intra-chain geometry allows us to simultaneously dock and design protein segments, while still targeting specific binding sites.

To better incorporate flexibility into our predictions, we provide only a coarse description of intra-chain geometry; presenting distance and angle information within a resolution of at least 2Å and 20^°^ respectively. On top of this, we attempt to approximate the unbound state of each training example, by applying Rosetta’s FastRelax protocol [31] to individual chains.

To validate our approach, we perform an extensive comparison against four other protein docking methods on unbound chains from Antibody Benchmark (AbBench)[32], and Docking Benchmark Version 5 [33]. We also show that DockGPT performs well in docking protein structures predicted by AlphaFold2 [34] with high success rates. Finally, we demonstrate how to extend DockGPT to perform simultaneous docking and *de novo* design by docking antibody-antigen partners while concurrently predicting both the sequence and structure of all heavy chain complementarity-determining regions (CDRs).

## 2 Related Work

### Geometric Deep Learning

The field of geometric deep learning is concerned with modeling data that has some underlying geometric relationships. Typically, this involves developing architectures that are *invariant* or *equivariant* with respect to the action of some symmetry group. Notable examples include the permutation equivariance of graph neural networks and the translation equivariance of convolutional neural networks.

Complementary to this work, several geometric deep learning methods tailored explicitly towards modeling symmetries of point clouds have recently been proposed [35–39]. These methods have helped facilitate significant improvements in protein-related molecular modeling tasks such as protein structure prediction [34, 40– 42], inverse folding [43–45], and *de novo* design [46–50].

### Traditional Methods for Protein Docking

Protein docking is traditionally performed in three steps: (1) sampling of candidate conformations, (2) score-based ranking of candidates, and (3) refinement of top-ranking complex structures. These algorithms primarily differ in either of the first two steps. Holding the position of the receptor fixed, each candidate conformation can be described by a 3-dimensional rotation and translation of the input ligand. Although the search space has relatively few degrees of freedom, the size of the effective candidate space can still total into the millions, even for small ligands [51]. In addition, the choice of score function usually induces a rugged energy landscape which is difficult to optimize over.

Within this paradigm, methods such as HDock [52, 53], PatchDock [54], ZDock [55], Attract [56], ClusPro [57], RosettaDock server [58], and Haddock [59], have been developed and made available for public access. Among these methods, PatchDock is one of the most widely used and computationally efficient. PatchDock avoids bruteforce search over transformation space by matching protein surface patches based on “shape complementarity.” Ligand transformations that align favorable patches bolster wide binding interfaces and avoid steric clashes resulting in favorable energy scores. HDock, ClusPro, and ZDock all make use of the Fast Fourier Transform algorithm to efficiently perform a global search on a 3D grid. The methods differ in how they post-process each candidate. ClusPro clusters candidates by root-mean-squared deviation and attempts to find a cluster with favorable energies. ZDock uses a combination of shape complementarity, electrostatics, and statistical potential terms for scoring. HDock, which was ranked as the number one docking server for multimeric protein structure prediction in the community-wide critical assessment of structure prediction 13 (CASP13-CAPRI) experiment in 2018, uses an iterative knowledge-based scoring function to discern the native complex. For a more complete review of traditional docking methods, available software, and accomplishments, we refer the reader to the compre-hensive reviews [60–62].

The majority of these methods incorporate backbone and side-chain flexibility only as a post-processing step, e.g. through molecular dynamics simulations. In order to incorporate inter-chain contacts or binding site residues, traditional docking methods typically alter their score function, or restrict search or results to ligand transformations matching this criteria. For example, ZDock allows users to specify “undesirble” residue contacts, and penalizes these interactions via the score function, and HDock applies post-processing to filter out predictions lacking target interactions.

### Machine Learning for Protein Docking

In the past, machine learning has been used outright or combined with physics-based methods for scoring docked complexes [63–65]. Recently, end-to-end machine learning methods EquiDock [30] and EquiBind [66] were proposed for protein docking and docking drug-like molecules. In particular, EquiDock makes use of an SE(3)-equivariant graph matching network to output a single rigid rotation and translation which, when applied to the ligand, places it in a docked position relative to the receptor. This is done by matching and aligning predicted *keypoints* which roughly correspond to the centroid of the binding interface. Although this method provides favorable theoretical guarantees, it does not perform well in practice. On top of this, the independent SE(3)-equivariant graph matching network and training procedure are relatively complicated. Training EquiDock requires solving an optimal transport plan which matches predicted interface key-points to ground truth positions, for each example. Custom loss functions are developed to back-propagate gradients through alignments and to penalize surface intersection. Moreover, it is unclear how to extend this method to account for conformational flexibility or more than two interacting chains. In contrast, our framework is conceptually very simple, utilizes standard architectural components and losses, allows for flexibility and is straightforward to extend to three or more chains.

### *De novo* Binder Design

Recently, there has been a spate of interest in *de novo* protein design using deep learning, especially the design of small protein binders. AlphaDesign [67] introduces a framework for *de novo* protein design which uses AlphaFold2 inside an optimizable design process, and [49] uses both AlphaFold2 and RosettaFold to improve the experimental success rate of their designs. Wang et al. [46] describe a method for *de novo* design of proteins harboring a desired binding or catalytic motif based on modifying the input and training of the RosettaFold network and augmenting the loss function. Here we show that our docking method can be easily extended to *de novo* design a protein that may bind a specific target site.

## 3 Methods

We overview our input representation, model training, loss, and architecture. Additional details can be found in Sections S1, S2 and S3

### Notation

We adopt the convention of using *x*_*i*_ and ***x***_*i*_ distinguish between a specific data point *x*_*i*_ and the list of data points (*x*_*i*_)^*i*=1..*n*^ indexed by *i*. We use **1**_*v*_ to denote the indicator function for a Boolean value v; evaluating to 1 if v is True and otherwise 0. A protein with *n* residues labeled 1..*n*, each with atom types ∈ 𝒜 *a*, is represented by its amino acid sequence ***s*** = *s*_1_, … *s*_*n*_, and atom coordinates 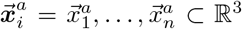. Each element *s*_*i*_ ∈ ***s*** can be any of the 20 naturally occurring amino acid types. To distinguish between multiple chains, we use *𝒞*_1_, …, *𝒞*_*k*_ ⊆ {1..*n*} to denote the partition of residue indices into chains 1..*k*.. We also use *C* (*i*) to denote the chain containing residue *i*, i.e. *𝒞* (*i*) ∈ {*𝒞*_1_, …, *𝒞*_*k*_}

### 3.1 Input Features

The input to our network is a complete graph 𝒢= (***x***_*i*_, ***e***_*ij*_) where 𝒱 consists of residue features *x*_*i*_ and ℰ consists of pair features *e*_*ij*_ between residues *i* and *j*. The bulk of our input features are generated independently, for each input chain. We refer to those features which do not depend on the input complex as *intra-chain features* and those which do as *inter-chain features*. In the interest of clarity, we first describe intra-chain features, which are independent of the protein complex being predicted.

#### 3.1.1 Intra-Chain Features

##### Residue Features

We generate residue features for each chain and join them by concatenating along the sequence dimension. The input feature *x*_*i*_ associated with residue *i* consists of four encodings:

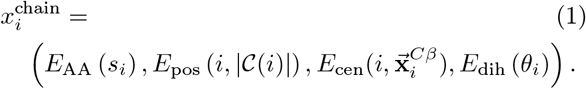

The first, *E*_AA_ (*s*), is a one-hot encoding of the amino acid type *s* using 20 bins for each naturally occurring amino acid. The next, *E*_pos_, encodes the residue relative sequence position into ten equal-width bins. As a proxy for estimating whether a residue is on the protein’s surface, we use a centrality encoding, *E*_cen_, which corresponds to the number of *Cβ* atoms in a ball of radius 10Å around the query residue. We encode this feature with six radial basis functions equally spaced between 6 and 40, and only consider residues in the same chain as the query atom. Last, *E*_dih_, encodes the angle *θ* ∈ [ − 180^°^, 180^°^] into 18 bins of width 20^°^. The input *θ*_*i*_ ∈ {*ϕ*_*i*_, *ψ*_*i*_} are the phi and psi backbone torsion angles of residue *i*. For residues before and after chain breaks, or at the N and C terminus of a chain, we set the phi and psi angles to 0.

##### Pair Features

Pair features are made up of low-resolution descriptions of pairwise distance and orientation and relative sequence information. The features for each chain are stacked to form a block-diagonal input matrix. A separate learned parameter is used to fill the missing off-diagonal entries. For a pair of residues *i* and *j*, in a common chain, the corresponding feature *e*_*ij*_ consists of three one-hot encodings

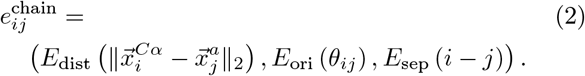

*E*_dist_ is an encoding of the distance *d* into six equal-width bins between 2Å and 16Å, with one extra bin added for distances greater than 16Å. We include distances between *Cα* and each atom type *a* ∈ { *N, Cα, C, Cβ*}. *E*_ori_, encodes the angle *θ* performed in the same manner as the backbone dihedral encoding for residue features. The input angles *θ*_*ij*_ ∈ {*ϕ*_*ij*_, *ψ*_*ij*_, *ω*_*ij*_} are pairwise residue orientations defined in [68]. Note that all pairwise distances and angles are known only within a resolution of at least 2Å and 20^°^, respectively. The last feature, *E*_sep_ (·), is a one-hot encoding of signed relative sequence separation into 32 classes, in the same manner as [69].

#### 3.1.2 Inter-Chain Features

We add three additional features to encode information about the target protein complex and PPIs.

##### Inter-Chain Interface (Residue)

*f*_*i*_ ∈ {0, 1} is an optional binary flag indicating whether the *Cα* atom of residue *i* is within 10Å of a *Cα* atom belonging to a residue in a different chain. This flag is 0 if this criteria does not hold.

##### Inter-Chain Contact (Pair)

*f*_*ij*_ ∈ {0, 1} indicates whether two residues in separate chains are in contact.

This occurs when the distance between 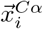 and 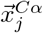 is less than 10Å. This flag is 0 if this criteria does not hold.

##### Relative Chain (Pair)

A one-hot encoding of the the relative chain index for residues *i* and *j* into three classes. Let *c*_*i*_, *c*_*j*_ ∈ {1, …, *k*} denote the chain index of residues *i* and *j*, then *f*_*ij*_ = OneHot (sign (*c*_*i*_ − *c*_*j*_)), where sign (*x*) ∈ {−1, 0, 1}.

The interface and contact flags provide context for residues on the binding interface for each chain; restricting the effective search space during inference. In realworld applications, knowledge of the binding interface may be limited or unknown. In light of this, we provide only a limited number of contacts or binding residues, chosen randomly for each training example. Specifically, for each input, we include no contacts or no residue flags, independently, with probability 1/2. This means that during training, the method sees 25% of examples without any interface or contact information, 50% with one or the other, and 25% with both features provided, on average. If interface features are included, we randomly subsample a number of interface residues *N*_int_ ∼ geom (1/5) to include, meaning five residues are selected on average. Similarly, we sub-sample *N*_con_ ∼ geom(1/3) inter-chain contacts when this feature is used, resulting in three provided contacts on average.

The relative chain encoding provides a way to distinguish between intra-chain and inter-chain pair features. By taking a signed difference, pair features *e*_*ij*_ and *e*_*ji*_ receive different encodings when *i* and *j* are in distinct chains. This not only discriminates the endpoints as belonging to different chains, but also breaks symmetry.

### 3.2 Deep Network Architecture and Training

We design a two-stage network making use of triangle multiplication, pair-biased attention, and invariant point attention (IPA). Our first module, which we refer to as the “structure encoder,” produces an invariant representation of the protein complex which is subsequently converted to 3D coordinates by the second module, the “structure decoder.” Our encoder uses pair-biased attention to update residue features, and triangle multiplication to update pair features. The decoder updates only residue features using IPA. We also make use of feature recycling during training and inference. We note that, although our architecture modifies or extends some components in AlphaFold2, the two architectures are functionally quite distinct. We do not make use of multiple sequence alignments (MSAs), templates, global attention, self-distillation, or other elements contributing to the success of AlphaFold2. In contrast, we hope to learn the principles governing protein binding from sequence and structure alone and develop a more specialized architecture to do so.

#### 3.2.1 Network Architecture

Here, we provide a general overview of our architectural components. A schematic overview of the architecture and loss can be found in Figure S1. Complete implementation details and more thorough descriptions of each submodule can be found in Section S1.

##### Structure Encoder Layer

Our encoder produces a joint representation of the input chains. Since interchain features are mostly missing from the input, we hypothesized that a network that updates pair features would facilitate more successful docking. Consequently, we chose to update pair features using incoming and outgoing triangle-multiplicative updates [34].

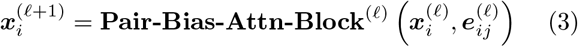

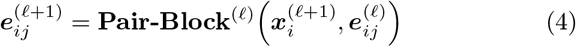

Each layer has two update blocks. The first block updates the residue features using multi-head attention with pair bias. The next block transforms the updated residue features into an update for the pair representation using a learned outer product, and then applies triangle multiplication and a shallow feed-forward network to the result.

##### Structure Decoder Layer

The decoder module converts the encoder representation to 3D Geometry. Since we do not make direct use of coordinate information in our input (although we show that this can be done for special cases in Section 3.2.1), we sought an invariant architecture specialized for coordinate prediction and ultimately settled on IPA.

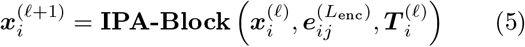

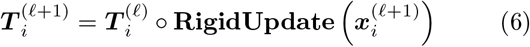

We use a total of six decoder layers, sharing the same weights for all six layers. We perform recycling during training and inference, allowing us to execute our model multiple times on the same example. This is done by embedding the previous iteration’s outputs in the next iteration’s inputs. Our best-performing model uses the same scheme as described in the AlphaFold2 implementation ([34], supplementary section 1.10). In concurrence with AlphaFold2 and OpenFold [70], we find that recycling significantly improves prediction quality while incurring only a constant increase in inference and training time. We experimented with recycling features from the structure decoder, rather than encoder. Since the decoder residue features encode plDDT information, we hypothesized that this information could better inform future iterations. This ablation and others are shown in Section S9.

##### Coordinate Prediction

In predicting residue-wise atom coordinates, we deviate from the strategy of AlphaFold2 and simply compute the local-frame coordinates for each atom using a learned linear projection. The coordinates for *Cα* are taken as the translation component of the per-residue predicted rigid transformation, and the remaining atom coordinates are predicted as

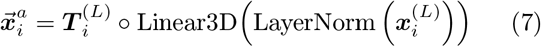

where *L* denotes the index of the last layer, and Linear3D is a learned projection into dimension |𝒜| × 3. Note that only one rigid transformation is used to produce all atom coordinates for a given residue.

##### Handling Coordinates as Input

Although we do not explicitly make use of coordinates in our docking model, for certain tasks, it may be important to incorporate this information as part of the input. This is especially salient in antibody loop design, where the framework region tends to remain mostly rigid upon binding. In Section S7 we show how to modify Equation (6) and Equation (7) to easily incorporate rigid, flexible, and missing coordinates as part of the input, while still maintaining SE(3)-Equivariance. We also provide empirical results for designing CDR loops with this modification in Section S8.

#### 3.2.2 Training

For general protein docking, model training is split into two stages. In the first stage, we pre-train on a mix of complexes and monomers, randomly selecting a monomer or a complex to train on with equal probability. This repeats for 5 epochs. The rationale for this decision is described in Section S3. Afterwards, monomers are removed, and we train exclusively on complexes. For all complexes in our training sets, we relaxed each individual chain using Rosetta’s FastRelax protocol [31] with all default settings (antibody heavy and light chains were relaxed jointly when applicable). For antibody-antigen docking results, we fine-tuned the model on a dataset consisting of only antibody-antigen complexes.

#### 3.2.3 Datasets

##### Single Chains

For pre-training, with single chains, we randomly sample chains from the publicly available BC40 dataset, consisting of roughly 37k chains filtered to 40% nonredundancy. Proteins with greater than 40% sequence similarity to any chain in our test datasets are removed.

##### General Protein Complexes

We use a subset of the publicly available Database of Interacting Protein Structures (DIPS)^1^ [71]. The training set is generated to exclude any complex that has any individual protein with over 30% sequence identity when aligned to any protein in the Docking Benchmark Version 5 test set (described in Section 3.3.2). We follow the training and validation splits for DIPS used in [72], with 33159 and 829 complexes, respectively.

##### Antibody-Antigen Complexes

For fine-tuning on antibody complexes, we use the publicly available Structural Antibody Database (SAbDab) which consists of 4994 antibody structures renumbered according to the Chothia numbering scheme [73–75]. Various papers from Chothia have conflicting definitions of heavy-chain CDRs ^2^. In light of this, we use the most recent definitions from [75]. We generate train and test splits based on antigen sequence similarity, filtering out examples where antigen chains have more than 40% sequence identity using mmseqs [76]. Before generating clusters, we removed all targets with overlap in our test sets, using the same criteria. We remark that no filtering is performed against antibodies. This results in roughly 3k complexes, for which we use a 8:1:1 split for training, validation, and testing.

#### 3.2.4 Loss

Our network is trained end-to-end with gradients coming from frame-aligned point error (FAPE), pairwise distance, per-residue lDDT (plDDT), and a few other auxiliary losses. We remark that our implementation of FAPE differs from that in AlphaFold2, as we use a different method for predicting coordinates. Other modifications were made in the clamping procedure of FAPE loss in order to facilitate faster convergence. A complete overview is provided in Section S2 and Figure S1.

### 3.3 Evaluation Criteria

To measure docking prediction quality, we report interface root-mean-square deviation (I-RMSD), ligand rootmean-square deviation (L-RMSD), DockQ score and DockQ success rate (SR) as reported by the DockQ algorithm^3^ [77]. DockQ score is a single continuous quality measure for protein docking models based on the Critical Assessment of PRedicted Interactions (CAPRI) community evaluation protocol. For antibody chains, we sometimes report CDR-RMSD which is calculated after superimposing the *Cα* atoms of the heavy and light chain framework regions using the Kabsch algorithm [78]. Finally, we sometimes include complex root-mean-square deviation (C-RMSD), which is derived by simultaneously superimposing all *Cα* atoms between two protein complexes. When assessing top-*k* performance, we take the best score over the top-*k* ranked predictions of each target.

When interface residues or contacts are specified, the information is randomly sampled from the native complex, and each method is run fifteen times for each target, each run with different random samples. For energy-based methods, outputs are ranked by predicted energy. For our method, we use predicted interface lDDT to rank each prediction. (see Section S4 for details).

#### 3.3.1 Docking Paradigms

In this paper, we are primarily concerned with predicting the bound conformation of a protein complex, given only unbound conformations of constituent chains. This is easily confused with *redocking*, the task of predicting a protein complex given *bound* conformations of each chain as input. Redocking is considerably easier than docking. For this task, traditional score-based methods are able to accurately predict most protein complexes. We verify this claim in Section S11.3, where we consider redocking antibody-antigen complexes.

When assessing docking performance, we sometimes condition on information about PPIs, such as interacting residues. Traditionally, amino acids are defined as interacting if any of their heavy atoms are within 6Å from one another. In this work, we used a more relaxed definition, where residues are defined as interacting if the distance between their *Cα* atoms is less than 10Å. This definition is more applicable to downstream protein design tasks, where knowledge of sequence or side-chain conformations may be missing or incomplete. In some cases, we provide the identity of select residues on the binding *interface* of a complex. In other settings, we provide *contacts*, which correspond to interacting residue pairs. We refer to the setting where neither interface residues nor contacts are specified as *blind* docking.

#### 3.3.2 Benchmarks

For each benchmark we include only receptor-ligand pairs having at least four contacts, and maximum chain-wise RMSD less than 10Å from the bound state. We note that some of the baselines might have used part of the DB5 test set in validating their models, and thus the scores may be optimistic. In addition to bound and unbound structures, we also include comparisons using receptor and ligand structures predicted by AlphaFold2 or AlphaFold-Multimer[41]. The same filtering criteria is applied to predicted structures.

##### Antibody Benchmark (Ab-Bench) [32]

A nonredundant set of 46 test cases for antibody-antigen docking and affinity prediction. This set contains both bound and unbound structures with diverse CDR-loop conformations between the bound and unbound states, ranging from ≤ 1Å to ≥ 4Å for CDR-H3. When AlphaFold-predicted structures are used as input, 26 test cases are used.

##### Docking Benchmark Version 5 (DB5) Test [33]

To the best of our knowledge, DB5 [33], which contains 253 structures, is the largest dataset containing both protein complexes and the unbound structures of their components. We use a total of 42 complexes from the DB5 test set which are held-out by the DIPS training split. For predicted structures, we also gathered 26 receptorligand pairs meeting the filtering criteria.

##### Rosetta Antibody Design (RAbD) [79]

A set of 46 κ and 14 λ antibody-antigen complexes. The benchmark contains antibodies with high CDR diversity and a wide range of length classes. All structures have experimental resolution ≤ 2.5Å, buried surface area in the antibody-antigen complex of ≥ 700^2^, and contacts with CDRs in both the light and heavy chain regions. These structures were used to assess the performance of docking algorithms in the bound input context, and results are given in Section S11.3.

## 4 Results

We compare DockGPT against ZDock [55], HDock [53], PatchDock [54], and EquiDock [30]. We downloaded their code and ran them locally. More details can be found in Section S10.

In addition to docking software, we provide a comparison with AlphaFold-Multimer [41] in Section S6, fig. S8, and table S1. We do not do so in the main text as the focus of this manuscript is protein docking and assessing the ability of docking programs to target specific binding sites. In general, current complex prediction algorithms such as AlphaFold-Multimer do not explicitly make use of binding site information, although it may be derived implicitly via multiple sequence alignments or templates. That is, they lack the ability to target specific binding modes, which further highlights the importance of effective docking methods.

### 4.1 Antibody Docking

We now compare methods on docking antibody-antigen unbound and predicted structures from the Antibody Benchmark dataset. As shown in Figure 2, for all but a few cases, our performance on docking AlphaFold2predicted structures roughly matches that on unbound inputs. In the interest of brevity, we report statistics for unbound inputs unless otherwise specified. Additional results and tables with RMSD and DockQ statistics are provided in Section S11.2. Results for docking RAbD bound chains are provided in Section S11.3.

**Figure 1:**
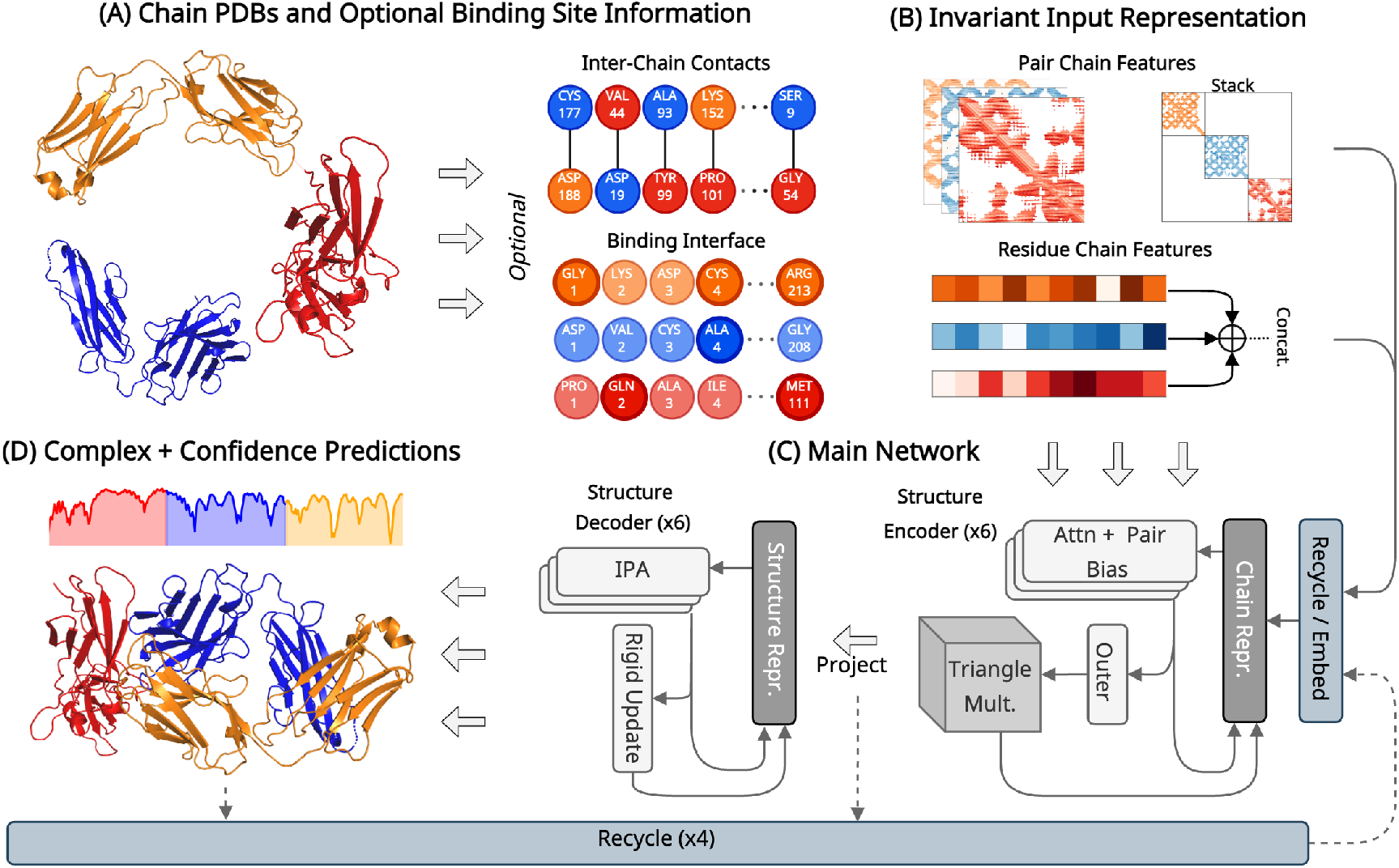
Approach Overview. (**A**) Unbound chain sequence and coordinates are given as input, and optionally, information regarding binding interface(s). (**B**) For each chain, an invariant representation of 3D geometry is constructed from quantities such as pairwise atom distances and orientations. If interface residues or contacts are provided, this information is added to the respective residue and pair features. Other features are discussed in Section 3.1. (**C**) The main network consists of two submodules. The *structure encoder* develops a joint representation of the input chains and the *structure decoder* infers the 3D geometry. (**D**) The output of the main network is the complex 3D-coordinates and per-residue confidence predictions. Steps (**C**) and (**D**) are repeated four times, with output residue, pair and distance features recycled from the previous iteration.

**Figure 2:**
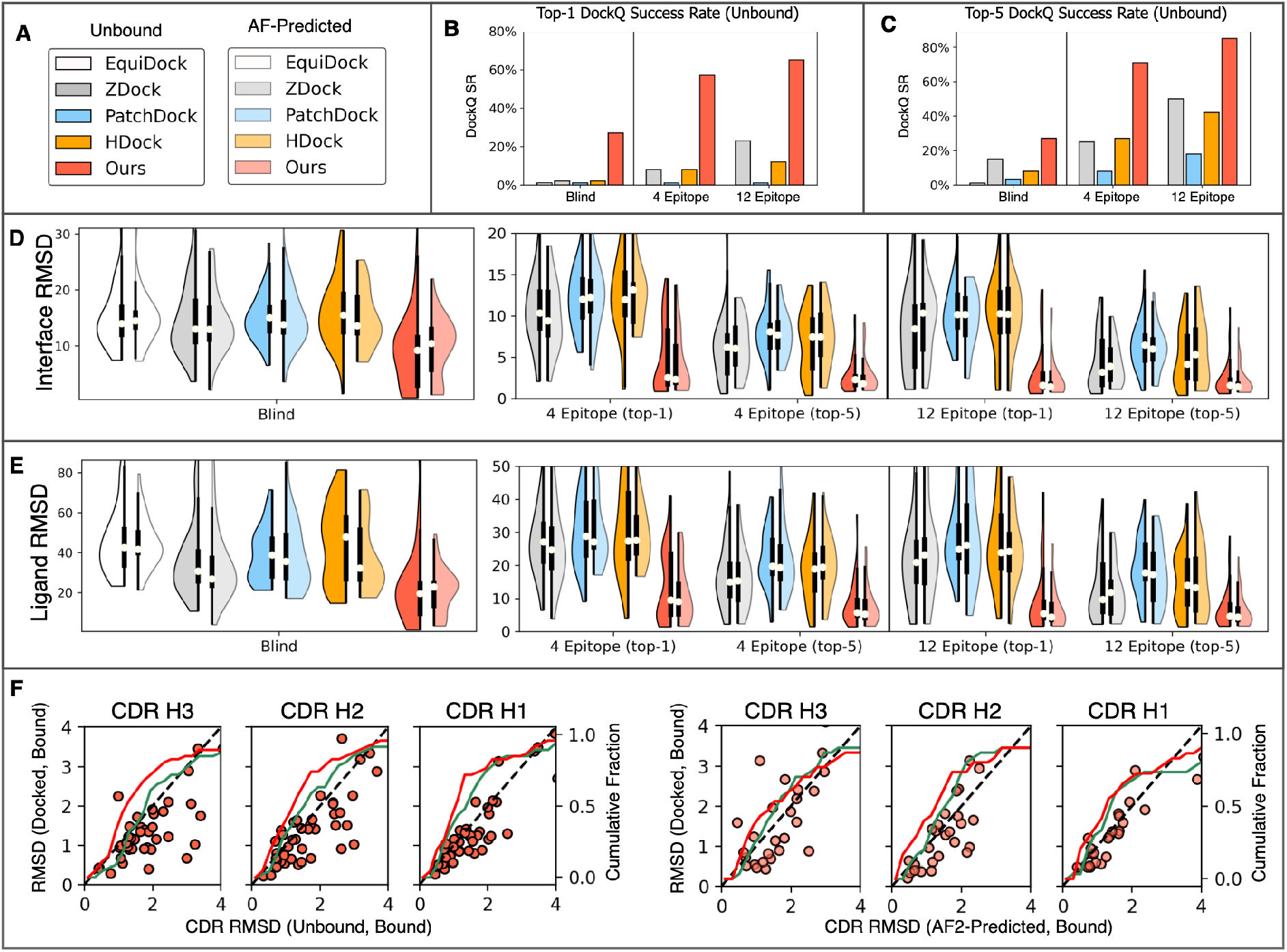
Results for Antibody Benchmark Predicted and Unbound Inputs. (**A**) Legends to distinguish between the five methods and the target type (predicted or unbound) in plots (B–E). (**B**) and (**C**) show top-1 and top-5 success rates for each method on unbound targets, with no epitope residues provided (blind) and 4 or 12 epitope residues provided. (**D**) Split violin plots showing interface RMSD distributions for docking unbound (left half) and predicted (right half) chains given 0, 4, or 12 epitope residues. Each violin plot marks the median value with a white dot, and shows the interquartile range with a bold vertical line. Both top-1 and top-5 distributions are shown when 4 and 12 epitope residues are provided. (**E**) Ligand RMSD distributions, in the same manner as (D). (**F**) Scatter plot of RMSD of heavy chain CDRs between our predicted and the ground truth (bound) complex structure. Here, the *x*-axis shows RMSD between the input (unbound or AF2-predicted) heavy chain CDRs and corresponding segments in the ground truth complex structure, and the y-axis shows RMSD between our predicted heavy chain CDRs and corresponding segments in the ground truth. Points below the y = *x* axis correspond to targets where the RMSD was lower for our predicted complex structures. The cumulative fraction of targets with CDR-RMSD less than the corresponding x value is also plotted on a secondary axis using a red line for our predictions, and a green line for unbound or predicted inputs. For these plots, we provided our method with 12 residues on the antigen epitope.

In Figure 2C and 2D, our method obtains top performance in blind docking (i.e., no interfacial contacts or residues are provided as input), with considerably lower interface and ligand RMSD values than others. This holds regardless of whether unbound or predicted structures are used as input. This carries over to DockQ success rate where our method exceeds 25% for both input types. Since our method is deterministic, we only make a single prediction in the blind setting, thus top-1 and top5 success rates are the same. In an attempt to improve blind-docking results, we developed a genetic algorithm which exploits our method’s ability to target different binding modes and predict docking quality. Details are given in Section S5 and examples are shown in Figure S6. This procedure increases both top-1 and top-5 success rates to 37.0% and 45.7% respectively.

For blind docking, traditional methods improve significantly when top-5 predictions are considered. ZDock’s top-1 predictions are successful for only one target, but this increases to 8 targets (17.4%) for top-5. Similarly, HDock improves median interface RMSD by roughly 5Å, from 15.6Å for top-1 to 10.8Å for top-5. EquiDock performs the worst of all five methods, with no DockQ successes for unbound or predicted targets. The method’s poor performance is likely a result of excessive steric clashes. On average, EquiDock has 581 backbone atom clashes between antibody and antigen chains. In contrast, our method does not produce more than 3 atom clashes for any target. Clash distributions for our method and EquiDock, along with some example predictions can be found in Figure S11.

Compared to the blind setting, performance for all methods improves significantly when information of the antigen binding interface (epitope) is included. When four epitope residues are given, we reduce top-1 median interface RMSD from 9.2Å to 3.1Å. Top-1 ligand RMSD decreases accordingly, from 19.5Å to 9.5Å. For traditional methods, the RMSD reduction is less dramatic. The best-performing traditional method, ZDock, decreases top-1 interface RMSD from 12.8Å for blind docking to 10.4Å when 4 epitope residues are given. Even when binding interfaces are accurately predicted, traditional methods often fail to orient protein backbones properly. When 12 epitope residues are provided, the lowerquartile interface RMSD of ZDock is 3.6Å, but the same quartile ligand RMSD is 14.4Å for top-1 predictions. On the other hand, our method obtains 1.2Å and 3.5Å RMSDs, respectively.

Parallel to blind docking performance, top-5 predictions of the traditional methods yield significantly higher DockQ success rates than top-1, when epitope residues are included. Furthermore, traditional methods see substantial improvements on all metrics when more epitope residues are provided. HDock and ZDock obtain top-5 DockQ success rates of 47.8% and 56.5% with 12 epitope residues, but only 30.4% and 28.3% with four residues. This is likely a side-effect of the post processing procedure, as increasing the number of epitope residues limits the size of the effective candidate space. In contrast, our method achieves a top-5 DockQ success rate of 71.7% with four epitope residues, increasing to 87.0% with 12. This suggests that our method has learned to use binding site information as more than just a search filter.

Considering the relationship between binding interface quality and prediction accuracy, in Figure 2E, we consider the heavy chain CDR-RMSD distribution of our docked antibody structures. Here, we see that CDR loop conformations predicted by our method are closer to the ground truth than that of the unbound or AF2-Predicted input. Predicted heavy-chain CDR conformations have median RMSD 1.55Å, 1.39 Å, and 1.81 Å compared to 1.82 Å, 1.67 Å, and 1.94Å for the unbound input. The outcome is similar starting from AF2-predicted input structures. This implies that our method goes beyond multidimensional scaling, and actually learns to incorporate backbone flexibility in its predictions.

Results for docking bound antibody-antigen structures are radically different than those shown in Figure 2. When blind-docking bound chains HDock and PatchDock achieve DockQ success rates of 79% and 25% respectively, for RAbD targets (see Figure S10). If we finetune and evaluate our model on bound antibody-antigen chains then the blind-docking success rate increases more than two-fold to 62%. This implies that important information about antibody-antigen binding interfaces is captured in the bound structures, and highlights the importance of comparing docking methods on benchmarks containing unbound structures. When training and analyzing on bound inputs, we still provide only a coarse description of geometry, and do not consider input sidechain conformations. We hypothesize that more finegrained featrues would significantly improve performance for bound targets. Interestingly, although EquiDock was trained on bound structures, the approach still underperforms on bound targets, with a success rate of 1.2%.

### 4.2 Results for DB5 Unbound and Predicted Targets

Results for DB5 targets are shown in Figure 3. Here, we focus mainly on performance when residue contacts are provided, but also consider providing a limited number of interface residues on one or more chains. We chose to provide at most *C* = 3 inter-chain contacts because, in theory, the number of rotational degrees of freedom for the ligand chain should be roughly max(0, 3 − *C*) if the contacts are well distributed. More results for DB5 predicted and unbound targets are shown in Figures 3, S4 and S5 and section S11.1, including tables with RMSD and DockQ statistics and performance on blind docking.

**Figure 3:**
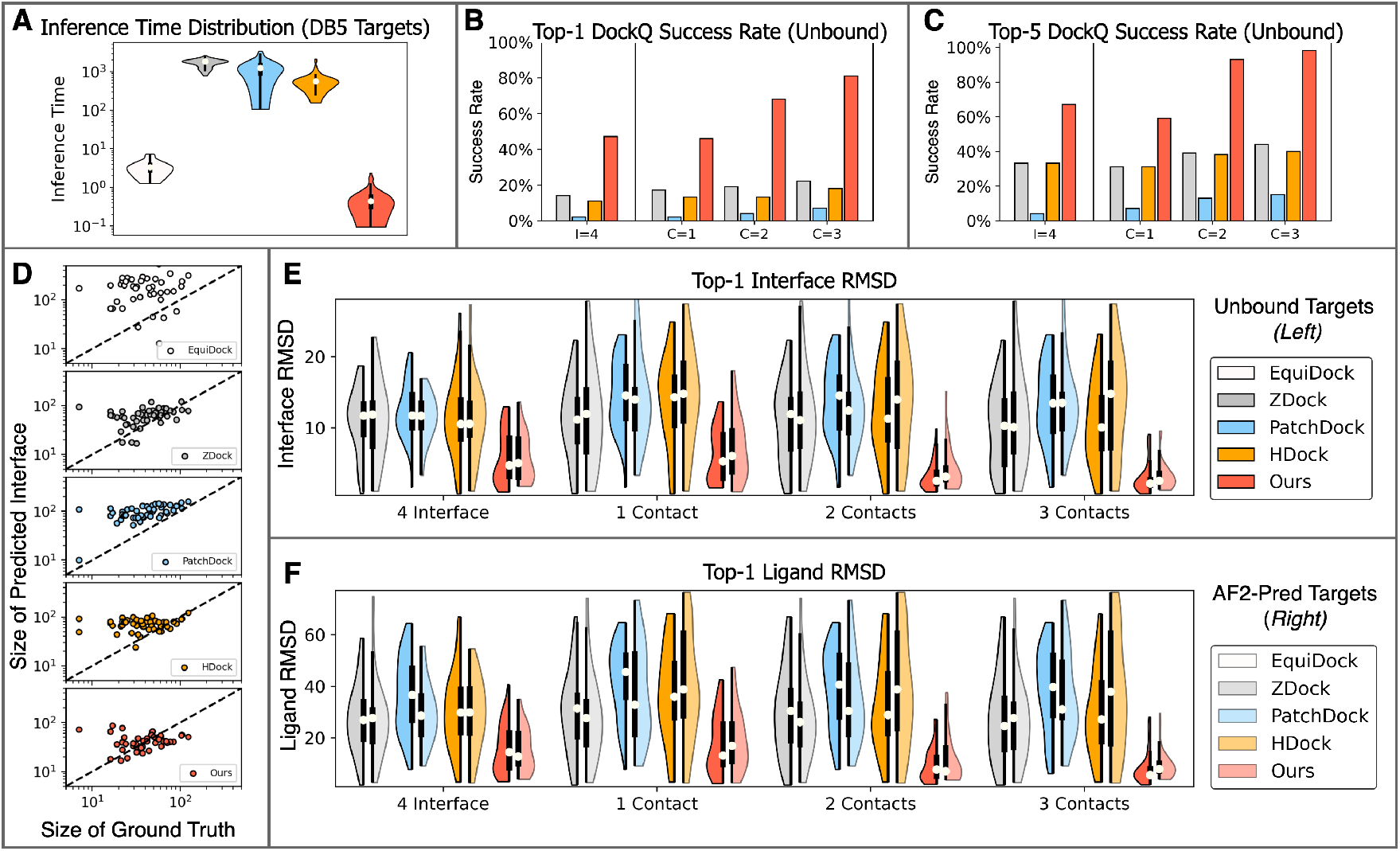
Results for DB5 with AlphaFold2 predicted and unbound monomer structures as input. (**A**) Per-method distribution of inference times for docking DB5 unbound targets. (**B**) Bar plot of Top-1 DockQ success rates for DB5 unbound targets. Each method was given one, two or three contacts (*C* = 1, 2, 3), or no contacts and four residues distributed over both the receptor and ligand binding interfaces (I = 4). (**C**) Bar plot of top-5 DockQ success rates, analogous to (B). (**D**) Scatter plot of the number of interacting residues in predicted complexes (y-axis) compared to that in the ground truth complex (*x*-axis). Blind docking predictions were made for all DB5 unbound targets, and interacting residues include only *Cα* atoms, with a cutoff distance of 10Å. (**E**) and (**F**) Split violin plots of Interface-RMSD and Ligand-RMSD distributions as in Figure 2. In (B,C,E,F) we exclude Equidock, because this method does not accept interface or contact information as input. Legends to distinguish between the five methods and target type are shown alongside RMSD distributions in (E) and (F).

For both unbound and AF2-predicted targets, supplying our model with a single contact generates better top-1 median RMSD scores than traditional methods supplied with up to three contacts. When one contact is given, DockGPT achieves a top-1 DockQ success rate of 45.2%, and 59.5% for top-5. In contrast, ZDock and HDock have less than 20% success for top-1, and 33.3% for top5. When 2 contacts are provided, DockGPT’s top-5 predictions are correct for all but three targets, and correct for all targets with 3 contacts, in terms of DockQ score. On the other hand, the success rate of traditional methods improves only moderately, with a maximum top-5 success rate of 45.2% for ZDock across all settings.

On top of performance, our method also achieves significantly faster inference times than others, averaging inference times more than three orders of magnitude faster than ZDock,HDock, and PatchDock, and approximately 6 times faster than EquiDock.

As shown in Figure 3D, blind docking predictions for methods EquiDock, HDock, and PatchDock tend to have large binding interfaces, even when there are few contacts in the ground truth complex. The tendency is most pronounced for EquiDock, which regularly predicts receptor-ligand poses with large surface overlap. On average, EquiDock, predicts a binding interface size that is 5.4 × larger than the ground truth. PatchDock, HDock average 2.9 × and 2.3 ×, that of the native complex, respectively. In contrast, ZDock and our method are the least biased, averaging 1.9× and 1.4×, respectively.

The tendency to predict large binding interfaces may be explained by considering the objective functions of each method. PatchDock explicitly rewards large binding interfaces and high shape complementarity. HDock and ZDock rank decoys by summing pairwise interfacial energy terms, and larger binding pockets may offer more potential for weak yet statistically favorable interactions. EquiDock, is trained to predict keypoints corresponding to the binding interface of each chain. It may be preferable from a loss perspective to predict keypoints near the chain’s center of mass when the binding interface is hard to discern. In theory, a chain’s center of mass offers a low-variance estimation of the true binding pocket. Finally, our method is trained with clamped FAPE loss and thus all predictions that deviate beyond the clamp value are equally “bad” in a loss sense.

An example of the behavior described in the previous paragraph is shown in Figure 4. Although EquiDock’s prediction is physically unrealistic, it still compares similarly to HDock in terms of interface and complex RMSD. EquiDock’s prediction has an interface RMSD of 14.8Å, and a complex RMSD of 17.2Å, whereas HDock obtains 20.5Å and 14.9Å respectively. It is also clear that HDock predicts a large binding interface for this target, even though the true binding interface is relatively small. This example also highlights the importance of assessing ligand RMSD in addition to complex and interface RMSD.

**Figure 4:**
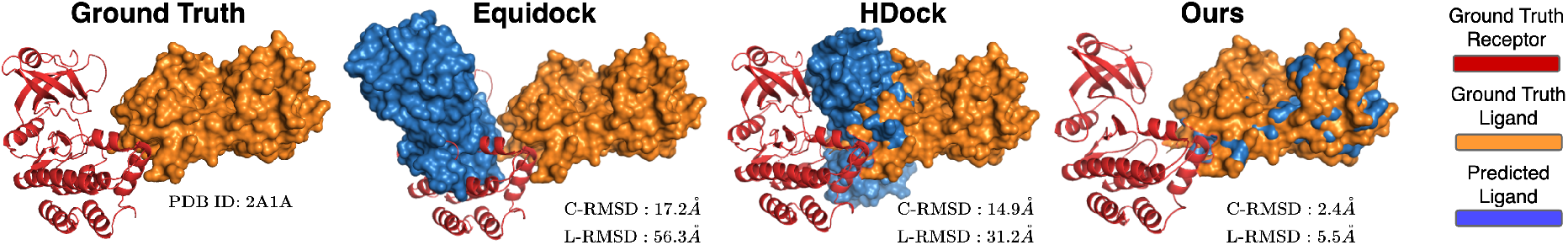
Predictions for one DB5 target with unbound structure as input. Ground-truth and docking predictions for PDB entry 2A1A. For each method, we show the surface of the predicted and ground truth ligand relative to the ground truth receptor. For this example, we selected traditional method HDock as it performed similarly or better than ZDock and PatchDock.

### 4.3 CDR-Loop Design

We now show how our model can be adapted to perform simultaneous docking, and sequence-structure co-design. For this task, we provide results for antibody CDR-loop generation, focusing on heavy chain CDRs H1-H3. We note that our method also designs light chain CDRs, but we omit this for brevity. In the remainder of this section, we briefly outline the modifications made to our approach and provide a comparison with other protein design frameworks tailored towards antibodies. More details and results can be found in Section S8.

#### Modifications to our approach

In order to perform joint imputation of sequence and complex structure, we retrained our model using data as described in Section 3.2.2, and all of same features as described in Section 3.1, except for residue degree centrality. We add one additional residue feature, which is a one-hot encoding of secondary structure using three classes for sheets, helices and loops. We encode all CDR residues as loops during inference, and do not apply masking to this feature during training. We found that the secondary structure encoding improved convergence when transitioning from pre-training to fine-tuning on antibody structures. During pre-training we masked linear segments of a randomly selected chain, sampling the segments based on proximity to the chain’s binding interface. The length of the masked segment is selected from a geometric distribution as geom 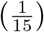. For each residue in the chosen segment, we replace the corresponding features with separate ⟨MASK⟩ parameters except for relative sequence position and relative sequence separation. To be clear, no inter-atom distance or orientation information is given to our deep learning model for masked residues.

#### Evaluation metrics and results

To generate our method’s results in Table 1, we provide four native antibody-antigen contacts, and produce five decoys per target. The decoy with the highest predicted interface plDDT is selected for the comparison.

**Table 1:** CDR-loop design. Performance of our method and four others on the task of predicting CDR H1-H3 loop conformation and sequence. For our method, “FT” denotes fine-tuning on antibody structures. The columns H1-H3 show the *Cα*-RMSD of heavy chain CDR H1-H3 between predicted and native structures. For our method, we also list the RMSD of the predicted and bound framework regions under column “Fr”. Perplexity (PPL) of sequence predictions for each CDR loop are shown in separate columns. Finally, overall perplexity and native sequence recovery across all loop regions is shown in the rightmost columns. We note that AR-GNN and RefineGNN predict sequence and structure for each CDR loop region separately, while conditioning on the native sequence and structure of the other CDR regions. This may result in slightly lower perplexity for these models.

| Method | Structure Prediction |  |  |  | Sequence Prediction |  |  |  |  |
| --- | --- | --- | --- | --- | --- | --- | --- | --- | --- |
|  | RMSD↓ |  |  |  | PPL↓ |  |  | NSR↑ | PPL↓ |
|  | H1 | H2 | H3 | Fr | H1 | H2 | H3 | H1-H3 |  |
| Ours | <u>1.11</u> | 1.02 | **1.88** | 0.72 | **4.46** | <u>6.71</u> | 10.68 | **39.7%** | 7.67 |
| CoordVAE [50] | **0.96** | <u>1.00</u> | <u>1.95</u> | – | – | – | – | – | – |
| Refine-GNN [48] | 1.18 | **0.87** | 2.50 | – | <u>6.09</u> | <u>6.58</u> | **8.38** | 35.4% | – |
| AR-GNN [48, 80] | 2.97 | 2.27 | 3.63 | – | 6.44 | 6.86 | <u>9.44</u> | 23.9% | – |
| LSTM [81, 82] | – | – | – | – | 6.79 | 7.21 | 9.70 | 22.5% | – |

Although our method receives only coarse information pertaining to antibody and antigen structures, we are still able to recover antibody framework regions with subAngstrom RMSD. Furthermore, none of the four other methods are capable of designing CDR loop regions in the presence of an antigen; for these methods the sequence and structure generation results in Table 1 are generated on bound heavy-light chains, with the bound antigen omitted. In contrast, our method simultaneously docks and designs all six heavy and light chain CDRloops and sequences simultaneously. Additional results and examples can be found in Section S8

## 5 Concluding Remarks

In this work, we developed DockGPT, a deep learning architecture for flexible protein docking with applications to *de novo* design of protein-binding proteins. Unlike other methods, our approach circumvents explicitly training on bound structures, and offers a natural approach to modeling conformational flexibility in complex prediction. By comparison across multiple benchmarks, we show that DockGPT meets or exceeds state of the art methods on rigorous quality metrics while also making better use of binding site information when it’s available. With significantly reduced inference times and explicit confidence estimates, we anticipate that our model will find further applications to machine-learning based virtual screening and *de novo* design platforms.

Despite our success, there are several limitations and extensions of our approach left open for future investigation. We use only a single atom type and threshold to supply our model with interface and contact information. Although it is straightforward to incorporate more finegrained binding site information, we did not explore this here. Parallel to this, supplying noisy or probabilistic binding site information could potentially improve performance and generalization. Although we do not provide explicit details in the main text, we remark that the current training procedure enables generation of diverse conformations by enumerating random contacts. We show in Section S5 how this can be used to rank and generate diverse binding modes, and ultimately improve blind docking. We suspect that this approach can be refined or extended to achieve even better performance. Although some of our deep network components were drawn from AlphaFold2, we do not incorporate any MSA information. We expect that MSA embeddings would be especially helpful in the blind docking setting. Finally, for *de novo* design tasks, we only analyzed our model on CDR loop design, and do not include estimates of binding affinity or free energy. Evaluation across more rigorous criteria and a broader range of design tasks must still be performed. We hope that future work will address some of these issues and develop this approach further.

## 6 Author Contributions and Acknowledgements

J.X. conceived and supervised the project and revised the manuscript. M.M. conceived and developed the machine learning algorithm, collected the results, built the datasets, and wrote the manuscript. M.M and J.X analyzed the results. The authors are thankful to members of the Xu group for helpful discussions. The authors also thank Xiaoyang Jing for generating AlphaFold2 predicted structures for our benchmarks.

## 7 Data Availability

Code is withheld until formal publication.

## Supplementary Information

### S1 Architecture and Hyperparamter Details

**Figure S1:**
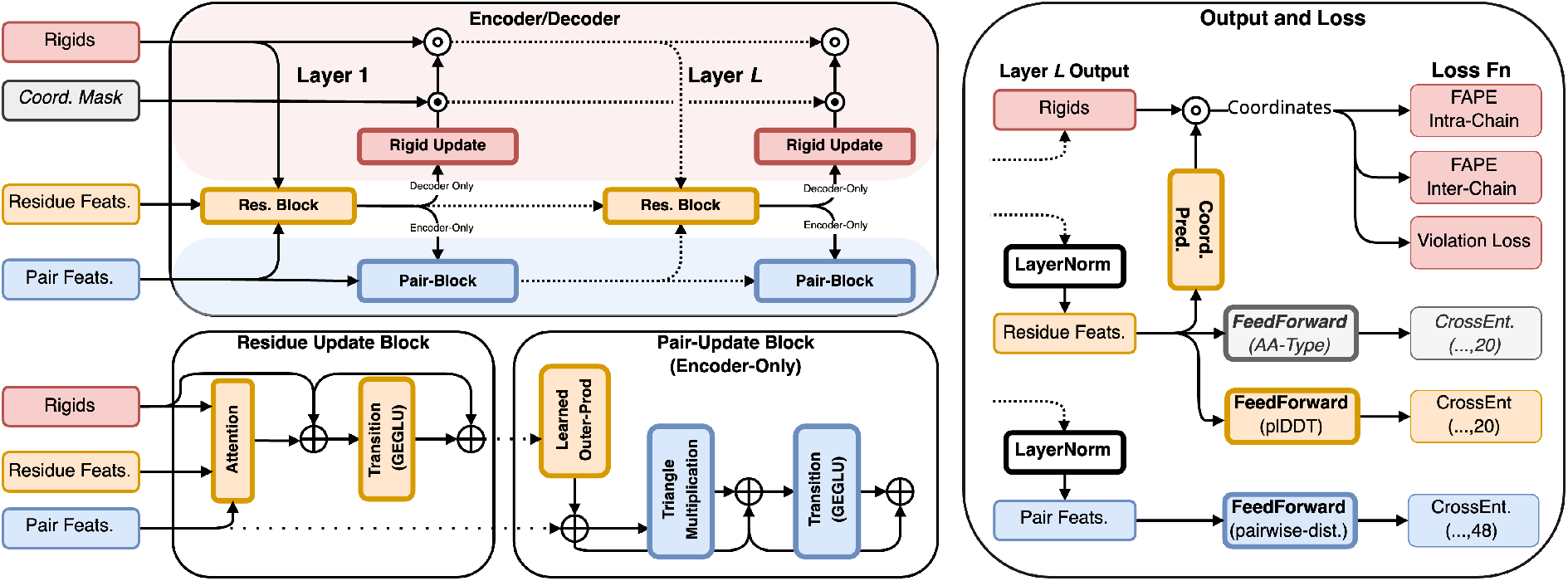
Architecture and Loss. Learnable modules are shown with bold text and bold borders. Modules operating on residue features are shown in orange and those operating on pair features are shown in blue. Modules making direct or indirect use of coordinates are shown in red. Optional input and modules for sequence and structure co-generation are shown in light gray. We use ⊕ to denote residual operations, ⊙ to denote element-wise multiplication, and ⊚ to denote the element-wise composition of two rigid transformations. Labeling of IPA and pair blocks with the index of their respective layer is omitted, as block weights may be shared across multiple layers. We highlight with a blue/red background those modules which are encoder/decoder specific (i.e. pair updates are omitted in the decoder, and rigids are omitted in the encoder). The structure encoder uses pair-biased multi-head attention for its residue update block, and the structure decoder used invariant point attention. Finally, layer normalization is applied, but not displayed here, except for in deriving the output.

Figure S1 shows a schematic overview of our model architecture and loss. We do not explicitly show our structure encoder module, since it differs only slightly from the structure decoder; the rigid update is removed, and IPA block is replaced with a pair-biased attention block. Though not displayed in the figure, we follow the Pre-LayerNorm scheme described in [83] where layer normalization [84] is placed inside the attention and transition residuals. In addition (pun intended), we use ReZero [85] for all residuals. Each feed-forward transition consists of one hidden layer having dimension four times that of the input dimension. For pointwise nonlinearity, we use gated GELU (GeGLU) based on success in other sequence modeling tasks [86, 87]. The Learned Outer-Prod module is nearly identical to that of the outer-product mean module described in [34], except we use a smaller intermediate dimension (*c* = 16 vs. *c* = 32), and skip the mean operation. The rigid update maps residue features to a per-residue rigid rotation and translation. This is implemented as a learned linear projection preceded by layer normalization. The composition of rigid transformations is implemented in the same manner as the backbone update in AlphaFold2 (see [34] Supp. Material, Algorithm 23). In some settings, the ability to incorporate prior coordinate information may be useful. Specifically, a subset of coordinates can be held fixed by replacing the corresponding rigid rotation and translation updates with the respective identity transformations (i.e. **I**_3_ and 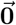). More details on this are provided in Section S7.

We use a hidden dimension of 256 for residue features and 128 for pair features in both submodules. All triangle multiplication updates use four heads of dimension 32 for queries and values. In the encoder submodule, we use eight attention heads of dimension 32 for each residue update block. Decoder IPA uses 12 heads per block, with dimension 16 for scalar features, and dimensions four and eight for point queries and values.

### S2 Loss Details

We use the same loss function for pre-training and fine tuning. Each term in the loss (shown in the equation below) is described in the remainder of this section.

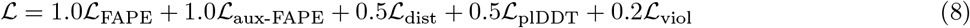

Similar to AlphaFold2, we also apply an averaged FAPE loss ℒ _aux-FAPE_, on the intermediate structures produced by the shared-weight layers of our decoder module. For the results in Sections 4.3 and S8, we add an additional term which is an averaged cross-entropy loss for amino acid identity, given weight 0.5

#### S2.1 Notation

We stick with the convention of using *x*_*i*_ and ***x***_*i*_ to distinguish between the individual data point *x*_*i*_ and the set of data points {*x*_*i*_ }^*i*=1..*n*^ indexed by 3D rigid transformations 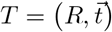 are represented by a rotation *R* ∈ *SO*(3), and translation 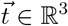. We use 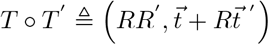 to denote the composition of two rigid transformations *T* and *T* ^*′*^. We use 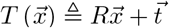 to denote the action of the rigid transformation on a vector 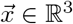. For notational convenience, and as a visual aid, we adopt the notation 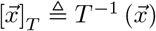 to denote the vector of coordinates *x* in the local frame defined by the rigid transformation *T*.

In the remainder of this section, we will use *n* to denote the number of input residues, and *𝒞*_1_, …, *𝒞*_*k*_ ⊆ {1..*n*} to denote the indices of residues in chains one and two respectively (i.e. *𝒞*_1_, …, *𝒞*_*k*_ is a partition of {1..*n*}). We assume that we have output residue features ***x***_*i*_, pair features ***e***_*ij*_, and predicted rigid transformations 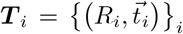 for each residue *i* ∈ {1..*n*}. We also assume predicted coordinates 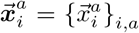 for each output atom type *a* ∈ 𝒜 derived as described in Section 3.2.1. When applicable, we use a superscript ∗ to distinguish between predicted and ground-truth data.

##### Per Residue lDDT

Residue output features are used to predict per-residue local distance difference test scores (plDDT). In defining the labels to evaluate on, there are two reasonable approaches. The first approach directly uses predicted coordinates,

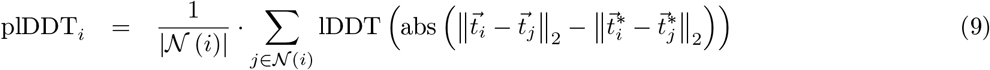

Where 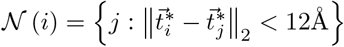, and

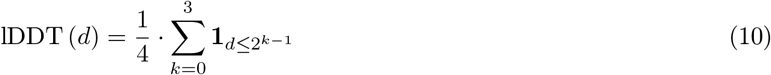

The alternative approach compares coordinates as they are seen in the predicted local frames of each residue. For this, we use predicted rigid transformations 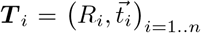 and true rigids 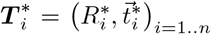 obtained from the native conformation to compute the local pLDDT score as:

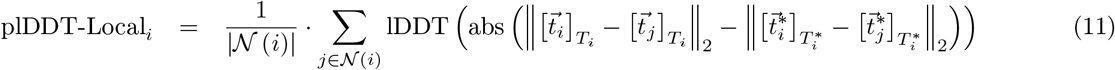

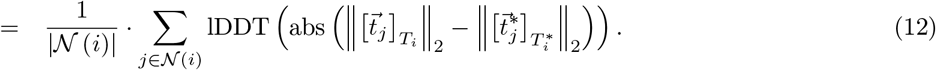

Ultimately, we use the standard plDDT to train our model. Although local-frame coordinates and distances are compared in each IPA head, we found that the local plDDT produces less accurate confidence estimates, and is also more difficult to optimize. Nevertheless, we include the alternative definition as it may be of interest to some readers.

To compute plDDT loss, we pass our output residue features ***x***_*i*_ through a shallow feedforward network with output representing 20 equal-width binned log likelihoods in the range [0, 1]. The predictions are compared with the ground-truth labels plDDT_*i*_ by cross entropy loss.

##### Pairwise Distance

We predict pairwise distances for four atom pairs (*Cα*, X), where X ∈ {*N, Cα, C, Cβ*} from 2-20Å using a bin width of 0.4Å. An extra bin is added for distances beyond 20Å. We do not separate inter and intra-chain atom pairs. Cross entropy loss is applied to compare the prediction to the ground truth.

##### Violation Loss

Unlike AlphaFold2, we predict only a single rigid transformation for each input residue. This means that intraresidue bond lengths and angles must be learned in the linear projection used to obtain predicted atom coordinates. We find that violation loss is very important for generating physically realistic conformations, and also for avoiding unfavorable steric interactions such as surface intersection. Here we use the same violation loss as defined in AlphaFold-Multimer; bond angle, bond length, and one-sided flat bottom steric penalty. We omit the “Center of Mass” loss [41, eq.1] as it had no empirical effect on performance.

##### FAPE

Here we describe a slight modification of the frame aligned point error (FAPE) loss described in [34, 41]. We reiterate that only a single rigid transformation is predicted for each residue, and thus rigid transformations for each output atom type cannot be directly compared.

Given predicted atom coordinates 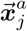 for each atom *a* ∈ 𝒜_*j*_ of residue *j*, we compute the per-residue FAPE, (pFAPE) for residue *i* as

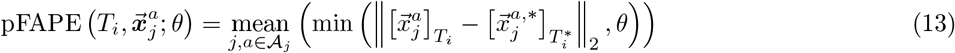

the FAPE loss over all residues is then

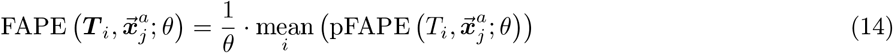

Our network employs two FAPE loss terms, each with equal weight. The first, FAPE_*intra*_ is is intra-chain FAPE which restricts the computation to pairwise relative coordinates within the same chain. The second is Inter-chain FAPE which applies the loss between atom coordinates in separate chains. Formally,

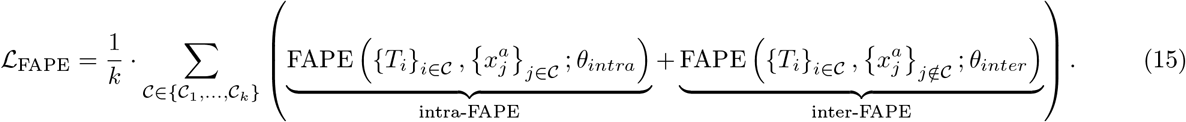

Following AlphaFold-Multimer, we use *θ*_*intra*_ = 10, and *θ*_*inter*_ = 30 with probability 0.9 and randomly set *θ* =∞ for each FAPE type with probability 0.1.

### S3 Training Details

All models were trained on 48Gb Nvidia RTX A6000 GPUs and optimized using Adam [88] with default parameters (*β*_1_ = 0.9, *β*_2_ = 0.999, ϵ = 10^−8^), with learning rate 10^−3^ during pre-training, and 5·10^−4^ afterwards. We apply per-example gradient clipping by global norm as described in [34, supplementary material, section 1.11.3], and scale the loss of each example by the log of the total number of residues to up-weight larger complexes. We validate our model every 500 mini-batches, using a minibatch size of 24. We train our model for at most 15 epochs, and apply early stopping with patience of eight validation steps. Since ReZero is used for residuals, we do not use any learning rate warm-up.

During the mixed monomer/multimer pre-training phase we crop complex chains so that the total number of residues does not exceed 500. We also remark that during the pre-training stage we append a binary flag to each residue and pair feature indicating whether the input corresponds to a single chain – in which case the chain should be treated as rigid. For general multimer training and antibody fine-tuning we place the encoder and decoder modules on separate GPUs and increase the crop size to 800 amino acids. Any complex containing a chain with more than 550 residues is removed from our training datasets. When cropping antibody-antigen complex chains, we randomly sample a contiguous subset of antigen residues so that the total number of resulting residues in 800. We follow the same strategy for general proteins, but choose a chain to crop at random. We note that no cropping was performed at inference time for any of the results in this paper.

#### Rationale for Single Chain Pre-Training

While developing this model, we first ran experiments to understand how well our architecture performed on multidimensional scaling tasks. For this, we sought to recover the *Cα* trace of protein chains given only distance and inter-residue orientation. We found that our deep model was able to recover the original *Cα*-trace with sub-angstrom RMSD using a 2Å resolution for distances, and 20^°^ resolution for angles after around 4k mini batches (approximately 1.5 epochs).

We attempted to apply the same model to rigid-docking, providing the same intra-chain information, but excluding all inter-chain features. In these experiments, the model struggled to reconstruct the conformations of the respective chains with reasonable accuracy, and showed a tendency to favor auxiliary loss terms such as intra-chain pairwise distance loss. This behavior persisted even after significantly more gradient updates (see Figure S2).

Considering this, we decided to separate FAPE loss into inter and intra-chain components, similar to what is done in [41], and pre-train our model on a 50-50 split of protein complexes and monomers. This resulted in significantly faster convergence in FAPE loss and far more accurate 3D-models. We remark that a single float (1 or 0) is appended to each residue and pair feature to indicate if the input is a complex or monomer.

**Figure S2:**
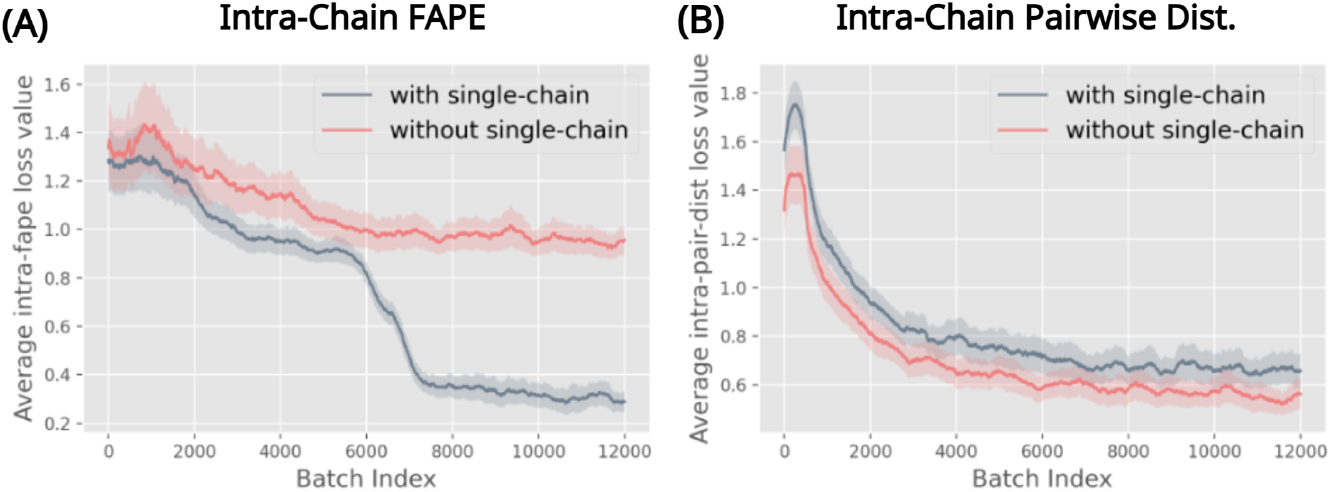
Training loss with and without monomer pre-training. (**A**) intra-chain FAPE loss (y-axis) and optimizer updates (x-axis). (**B**) Intra-chain pairwise distance loss (y-axis) and optimizer updates (x-axis).

### S4 Decoy Ranking with Predicted lDDT

In Section 4 we mention that decoys for each docking target are ranked by predicted interface lDDT (I-plDDT). We now describe this procedure in more detail. We define the predicted binding interface as the set of residues having at least one predicted inter-chain contact; a pair of residues from distinct chains, with *Cα* distance between predicted coordinates less than 10Å. The true (actual) binding interface is defined analogously with respect to the ground truth complex. To rank decoys for a given target, we take an average of the per-residue lDDT as predicted for those residues on the predicted binding interface. The plDDT score for a given residue is taken as an expectation with respect to the predicted logits. Similarly, the predicted lDDT for a decoy is defined as the average over predicted plDDT for all residues in the complex.

**Figure S3:**
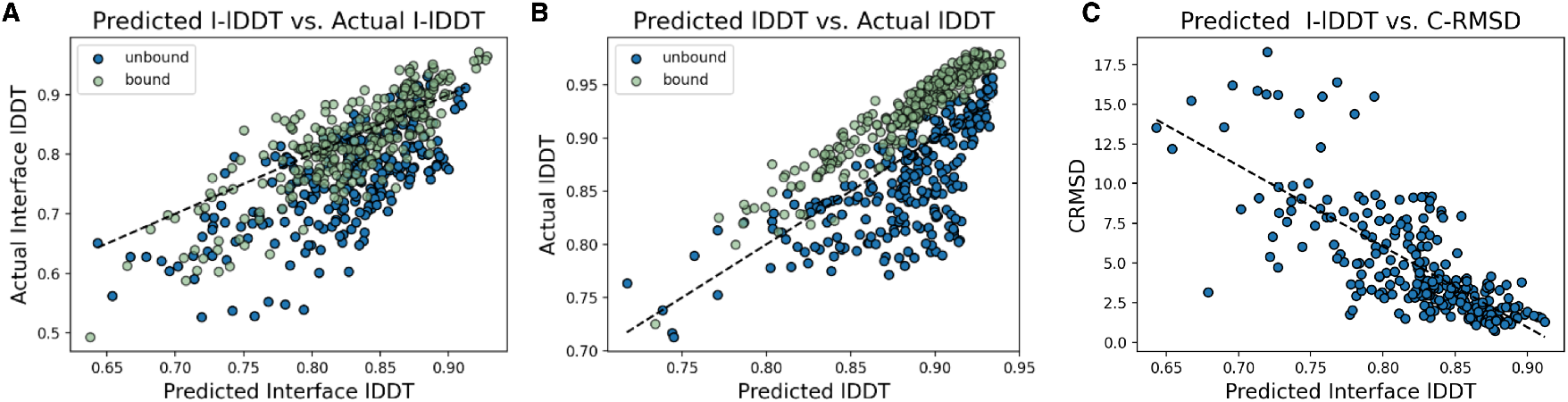
Analysis of lDDT predictions. Each plot shows results for predictions made on DB5 bound or unbound input chains, providing the model four interface residues and four contacts sampled at random from the ground truth complex. Each dot represents a decoy generated from bound or unbound input chains. A total of five decoys were generated for each target. Correlation coefficients for predictions derived from unbound and bound targets are denoted with ρ_*u*_ and ρ_*b*_ respectively. (**A**) scatter plot of predicted lDDT (x-axis) for the predicted binding interface against actual lDDT (y-axis) for the ground truth binding interface. Unbound targets are shown in blue (ρ_*u*_ = 0.69) and bound targets are shown in green (ρ_*b*_ = 0.83). We remark that the predicted and actual interfaces may differ. (**B**) Scatter plot of predicted lDDT (x-axis) and actual lDDT (y-axis) for bound and unbound targets (ρ_*u*_ = 0.70), ρ_*b*_ = 0.95). (**C**) Scatter plot of predicted lDDT using the predicted binding interface against the complex RMSD of the predicted structure (ρ_*u*_ = −0.74).

Figure S3 shows scatter plots of predicted lDDT and predicted I-lDDT for DB5 bound and unbound targets. In plot (C), we find a strong correlation between I-plDDT and complex RMSD for unbound targets, suggesting that this quantity is effective for ranking decoy structures. We explore this further in Figure S4, which compares the complex (A) and interface (B) RMSD distributions of decoys selected by plDDT (orange) and the same distributions computed over all decoys (blue). In this figure, we again generate five decoys per target, and assess across 12 binding site settings, varying the number of provided contacts or interface residues in each setting. Mean and median RMSD scores for selected decoys are lower across all binding site contexts. RMSD distributions of decoys selected by interface plDDT are also consistently more concentrated at lower values.

**Figure S4:**
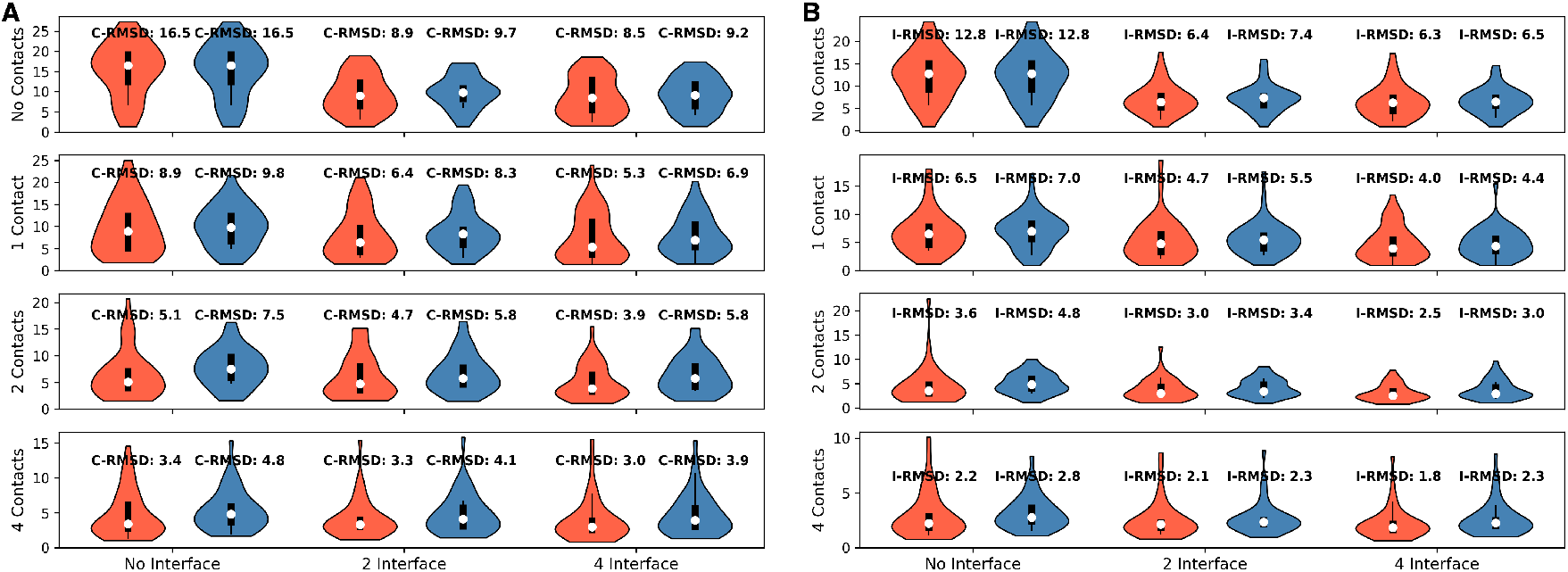
Selection overview for DB5 unbound targets (without recycling iterations). For this experiment we generate five decoys for each target using a reduced model (no side chain prediction, no recycling). Each row/column corresponds a number of provided contacts/ interface residues. This information is derived as a random sample from the native conformation. For each violin plot, we compare the complex RMSD (C-RMSD, (**A**)) or Interface RMSD (I-RMSD, (**B**)) of all predictions (blue) against the prediction for each target having highest predicted interface plDDT (orange). We remark that results in the two plots use only *Cα* atoms to compute each RMSD type, and as such, may differ slightly from the results reported in other sections.

**Figure S5:**
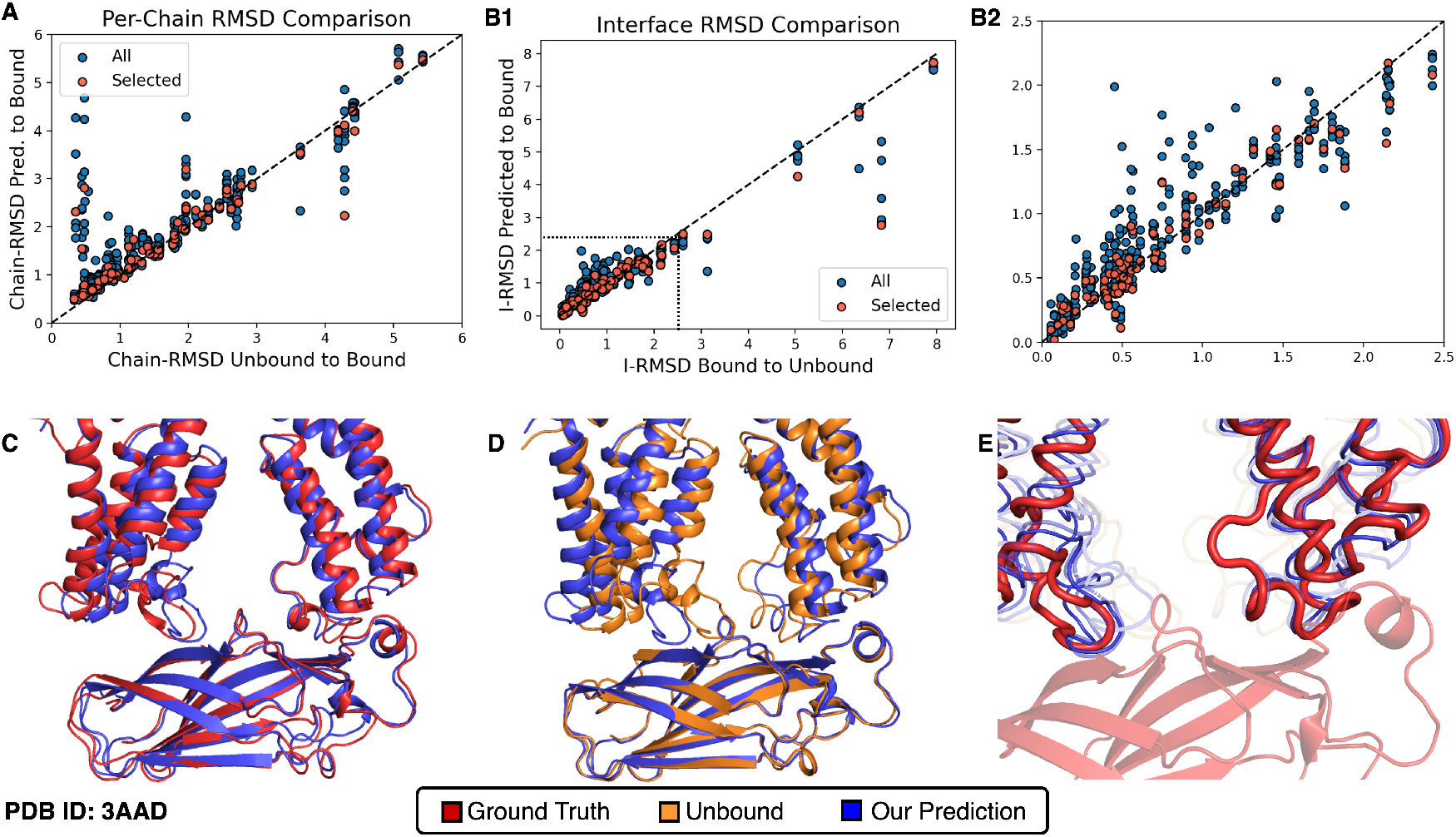
Examination of conformational flexibility for DB5 unbound targets. As in Figure S3, we generate 5 predictions per DB5 target, using unbound chains as input to our model. For each prediction, we provide our model with three contacts sampled at random. (**A**) Scatter plot of receptor/ligand chain-wise RMSD between bound and unbound chains (*x*-axis) against predicted and bound chains (y-axis). Red dots show the decoy with highest predicted interface pLDDT for each target. (**B1**) Shows the interface RMSD in the same manner as (A). (**B2**) zooms in on the 0-2.5Å range of (B1). (**C**–**E**) Cartoon representations of our prediction, bound, and unbound chains for DB5 target 3AAD. (**C**) Our top-ranked prediction for DB5 target 3AAD using unbound chains as input is shown in in blue, and the bound conformation is shown in red. (**D**) Cartoon representations of our top-ranked prediction (blue) and unbound chains (orange) for target 3AAD. For this image, unbound chains are optimally aligned to respective bound chains using a chain-wise Kabsch alignment. (**E**) Our model’s top-3 ranked predictions for 3AAD, colored by predicted interface lDDT. Lower transparency is used to denote lower predicted interface-LDDT. For this target, the RMSD between bound and unbound receptor chains (top, helices) is 4.18Å, and 2.05 Å for the ligand chain (bottom, sheets). The interface RMSD is ≈ 6.8Å when bound and unbound chains are optimally aligned. Our top ranking prediction obtains an interface RMSD of 2.6Å.

Last, we consider our model’s ability to predict conformation changes upon binding. In Figure S5(A,B) we see that the chain-wise RMSD between predicted and unbound structures is similar for all but a handful of targets. In terms of interface RMSD, predicted structures are slightly more similar to that of the bound conformation, especially when there are larger discrepancies in the interface of aligned bound and unbound structures.

Unfortunately, the conformation similarity between DB5 bound and unbound structures is relatively high, and more diverse structures should be examined before drawing conclusions from these results. Nevertheless, in Figure S5 (C and D) we consider a case study on PDB entry 3AAD, where our model predicts a conformation diverging significantly from the unbound state. For this target, our model with highest predicted interface lDDT has interface RMSD 2.6Å, where as an optimal alignment mapping the unbound chains to the bound complex has interface RMSD 6.8Å. Moreover, our model predicts a conformation for the helical receptor chain that is only 2.2Å from that of the bound conformation; compared to 4.2 for unbound-bound conformation. We remark that the maximum sequence identity between target 3AAD and any training example is only 9%.

### S5 Genetic Algorithm for Protein-Protein Docking

Although our method is deterministic, sampling can still be performed by providing different subsets of interchain contacts of binding interface residues for the same example. To sample conformations in the absence of interfacial residue and contact information, we use a genetic algorithm to guide complex predictions towards high confidence binding modes. Any genetic algorithm consists of three main components: (1) a genetic representation of the solution domain, (2) a “fitness” function to assess population quality, (3) a mutation function which alters representations, and (4) a crossover function which combines two representations. Given an initial population of solution candidates, the algorithm then proceeds to produce new “generations” by assessing the fitness of each candidate and stochastically selecting those with favorable fitness to combine or mutate. We run this procedure for a total of 10 generations, using an initial population size of 50, and subsequent population sizes of 25. We describe each component of our algorithm below.

#### Solution Representation

Solutions are represented as a binary vector of interface residues. The length of this vector is L_*rec*_ + L_*lig*_ where L_*rec*_ is the length of the receptor chain, and L_*lig*_ is the length of the ligand chain. Each position of the vector corresponds to a residue in one of the chains, and a one at position *i* is meant to indicate that this residue *i* is part of the binding interface.

#### Initial Population

To generate initial candidates 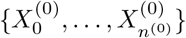, we randomly sample a single residue on the surface of receptor and ligand chains, and provide these two residues as the “interface-residue” feature. For antibodies, we restrict the sampling to residues in CDR H1-3 loops. Random surface residues are chosen by scaling a 3-dimensional Gaussian (direction), to the maximum distance between any two residues in the protein, and then choosing the residue closest to this point.

#### Fitness Function

To evaluate the fitness of each candidate, we use the candidate solution as the binding interface feature for our method, and then compute a function of predicted interface-pLDDT on the output. We choose *f*(X, t) = exp[t · (I-pLDDT(X))] where t is a scaling parameter (chosen ad hoc as one plus the index of the current iteration).

#### Mutation Function

Given a set of solution candidates, 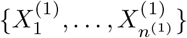 and corresponding structures gener-ated at time t, we select a subset of *n* = *n*^(*t*+1)^ with replacement according to the fitness function *f*(·, t + 1), and randomly sub-sample six residues on the predicted binding interface. We choose to sample a fixed number here because we empirically found that predicted interface lDDT scores have a modest correlation with the number of interface residues provided as input.

**Figure S6:**
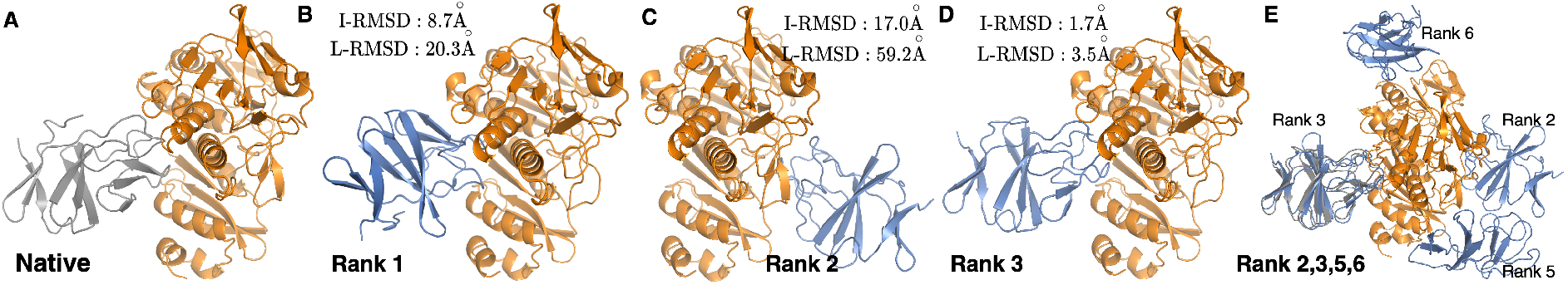
Genetic Algorithm Explores Diverse Binding Modes. Ground truth and example predictions from our genetic algorithm for DB5 target 2YVJ. In all sub-figures, the ground truth receptor is shown in orange, the bound ligand is shown in gray, and our predictions are shown in blue. (**A**) Bound complex of DB5 Target 2YVJ. (**B**–**D**) the top three ranked predictions using our genetic algorithm. (**E**) Rank 2, 3, 5, and 6 predictions from our genetic algorithm. Rank 1 and rank 4 predictions are omitted for visual clarity, as they clash with some other predictions. The bound ligand is also shown in gray. Although our method fails to generate an accurate top-1 prediction, our third ranked prediction successfully docks to the same interfacial region.

### S6 Comparison to AlphaFold-Multimer

We compare our method with AlphaFold-Multimer in the blind docking setting on DB5 and Ab-Bench benchmarks described in Section 3.3.2. In addition to comparing the two methods directly, we also include a hybrid approach (Ours + AF). For this approach, we provide our method with up to three randomly sampled residues from antibody-antigen binding interfaces predicted by AlphaFold-Multimer. No information of native complexes is used for our method. We generated 100 decoys for each target, and selected the decoy with highest predicted interface lDDT as our final prediction (selection as described in Section S4). The results are shown in Table S1.

**Figure S7:**
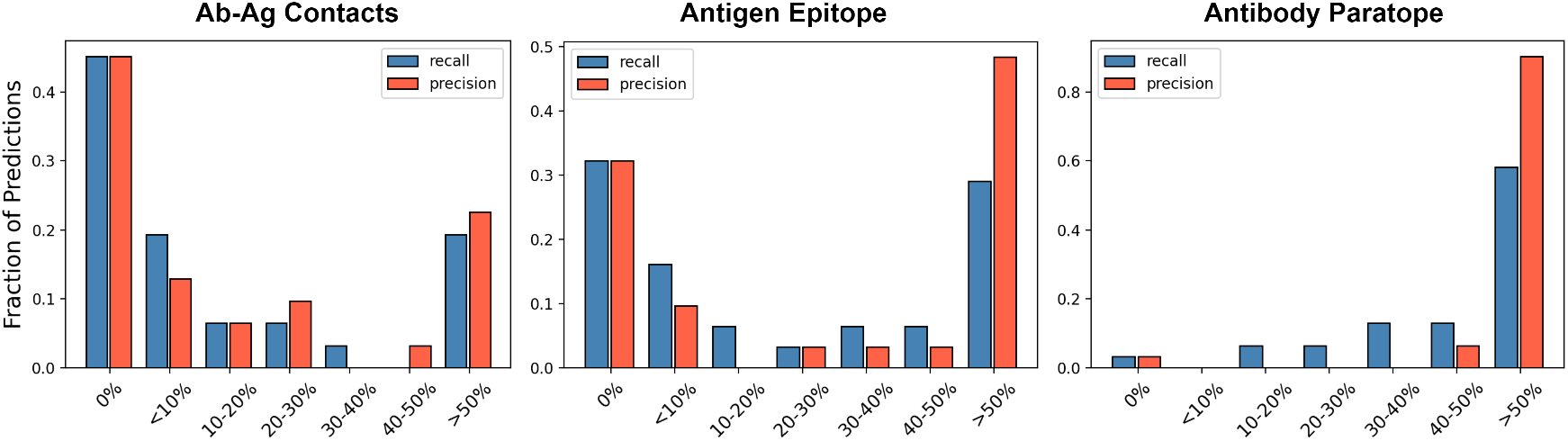
Binding site precision and recall for AlphaFold-Multimer on Ab-Bench targets. Histograms of binding site precision and recall for AlphaFold-Multimer predicted structures on Ab-Bench targets. Recovered contacts, antigen binding interface (epitope) and antibody binding interface (paratope) is shown from left to right.

Motivating the hybrid approach, we analyzed binding site information extracted from AlphaFold-Multimer predictions (Figure S7). As expected, AlphaFold-Multimer recovers the antibody paratope with high precision. Perhaps more surprising, we see that at least part of the antigen epitope is recovered with relatively high precision, but lower recall. Noticing this, we conjectured that our results may be improved by sampling a limited number of predicted binding modes and ranking predictions.

**Table S1:** Comparison of Our Method and AlphaFold-Multimer on Two Docking Benchmarks. Results for AlphaFold-Multimer (AFM), our method (ours), our method with genetic algorithm (Ours+GA), and our method using AlphaFold-Multimer predicted interfaces (Ours+AFM) for Ab-Bench and DB5 benchmarks. AlphaFold-Multimer outperforms our method on blind docking general protein targets from DB5. Our method does not make use of MSA information, which is especially important for general proteins where binding interfaces are harder to discern. For antibody complexes, the paratope is limited to CDR loops and our method has an easier time predicting the complex.

|  | Top-1 |  |  |  |  |  |  | Top-5 |  |  |  |  |  |  |
| --- | --- | --- | --- | --- | --- | --- | --- | --- | --- | --- | --- | --- | --- | --- |
|  | DockQ↑ | I-RMSD↓ |  |  | L-RMSD↓ |  |  | DockQ↑ | I-RMSD↓ |  |  | L-RMSD↓ |  |  |
|  | SR (%) | 25<sup>th</sup> | 50<sup>th</sup> | 75<sup>th</sup> | 25<sup>th</sup> | 50<sup>th</sup> | 75<sup>th</sup> | SR (%) | 25<sup>th</sup> | 50<sup>th</sup> | 75<sup>th</sup> | 25<sup>th</sup> | 50<sup>th</sup> | 75<sup>th</sup> |
| Antibody Benchmark |  |  |  |  |  |  |  |  |  |  |  |  |  |  |
| AF-Mult. | 28.3% | <u>1.9</u> | 9.3 | 14.7 | 12.2 | 22.6 | 36.0 | 34.8% | 1.8 | 5.8 | 13.1 | 9.2 | 18.3 | 26.4 |
| Ours | 26.1% | 2.5 | <u>9.2</u> | **12.1** | 8.2 | <u>19.5</u> | <u>25.4</u> | - | - | - | - | - | - | - |
| Ours+GA | **37.0%** | **1.8** | **8.3** | <u>12.4</u> | **5.5** | **19.2** | <u>26.4</u> | **45.7%** | **1.7** | **4.0** | **7.3** | **4.9** | **11.5** | **19.8** |
| Ours+AFM | 28.3% | <u>1.9</u> | 10.1 | 13.3 | <u>5.7</u> | 20.1 | 27.8 | 37.0% | **1.7** | <u>4.4</u> | <u>8.9</u> | <u>5.4</u> | <u>11.3</u> | <u>19.0</u> |
| Docking Benchmark Version 5.5 |  |  |  |  |  |  |  |  |  |  |  |  |  |  |
| AFM | **50.0%** | **0.9** | <u>7.9</u> | <u>16.4</u> | **2.8** | <u>19.6</u> | <u>35.2</u> | <u>50%</u> | **0.9** | **4.7** | <u>13.0</u> | **2.6** | **12.2** | <u>30.2</u> |
| Ours | 7.1% | 8.9 | 13.3 | 17.4 | 24.2 | 35.4 | 49.5 | - | - | - | - | - | - | - |
| Ours+GA | 9.5% | 9.7 | 14.0 | 17.5 | 23.1 | 33.4 | 47.5 | 16.7% | 5.4 | 8.8 | 13.6 | 12.9 | 20.7 | 34.3 |
| Ours+AFM | <u>42.8%</u> | <u>2.7</u> | **5.7** | **14.3** | <u>6.6</u> | **17.4** | **28.7** | **52.4%** | <u>2.0</u> | <u>5.1</u> | **12.7** | <u>5.2</u> | <u>12.4</u> | **24.5** |

Our blind docking (i.e., our deep learning plus our genetic algorithm) greatly outperforms AF-Multimer on antigen-antibody complex structure prediction without using any binding site information. But on general protein targets, our method performs poorly. Adding AF-Multimer predicted interface or contact information significantly improves prediction quality since this indirectly makes use of MSA information. We hypothesize that directly including MSA information could significantly improve prediction quality for general proteins, especially in conjunction with our genetic algorithm, as model confidence predictions correlate strongly with predicted interface plDDT, but we leave this study for future work.

Recently, Yin et al. [89] benchmarked AlphaFold-Multimer and other docking programs on antibody-antigen and general protein targets using the sequences or structures of unbound chains. This study found that AlphaFoldMultimer performs very poorly for antibodies, successfully predicting only 11% of targets. In their study, the authors identified sequence and structural features associated with lack of AlphaFold success and attribute the performance gap to lack of co-evolutionary signal. For antibody-antigen complexes, they found that the success rate of AlphaFold-Multimer was not much different when the model was given only templates, and no MSA information. In this setting, AlphaFold-Multimer is similar to our model. We hypothesize that our performance improvement for antibody-antigen targets comes from (1) fine-tuning and (2) no MSA inputs. Since we do not train with MSA information, our model is forced to learn sequence and structural features which facilitate good binding modes. This is particularly useful for immunoglobulin targets, as antibody-antigen interfaces are less likely to have co-evolving sequences available for MSA generation [89].

**Figure S8:**
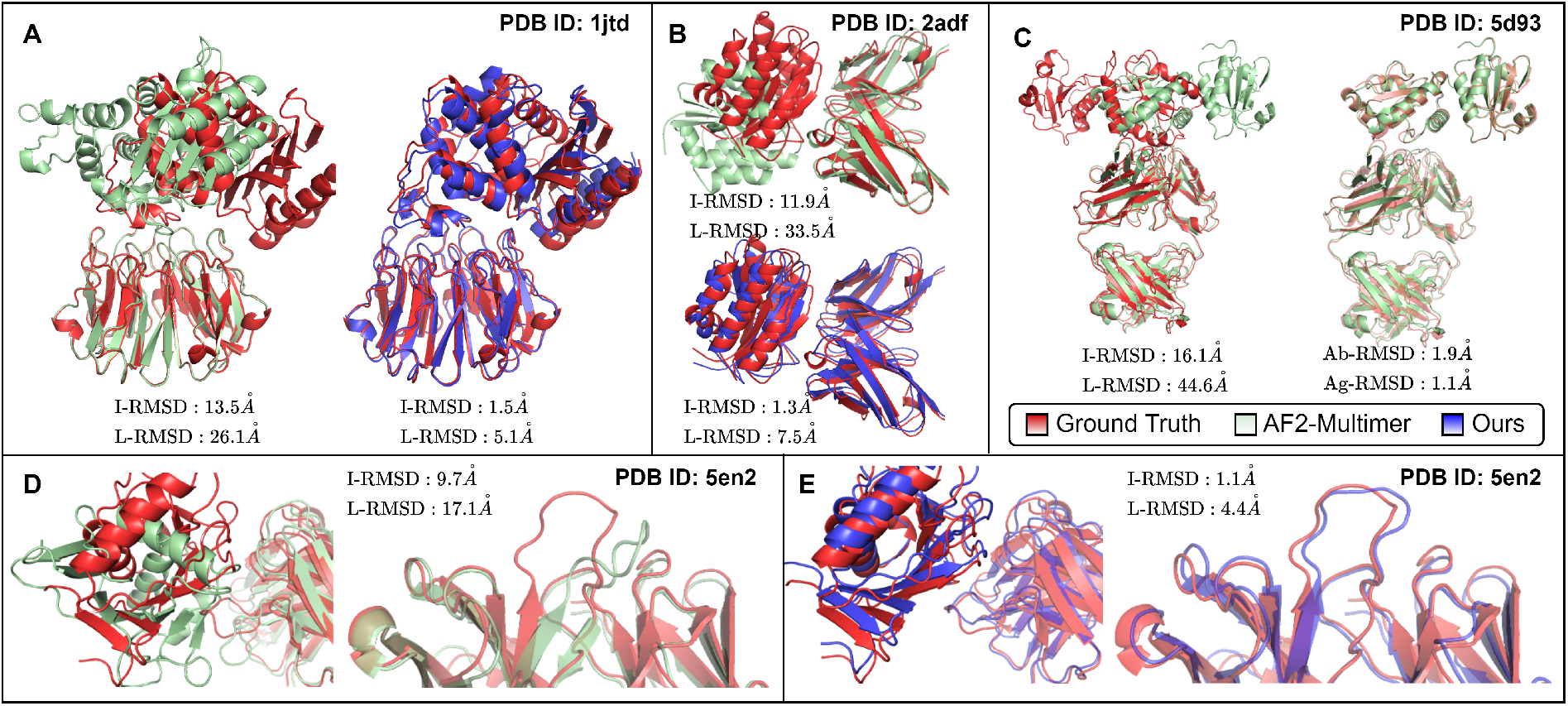
Comparison of Structure Predictions Between Our Method and AlphaFold-Multimer. In this figure, all predictions from our model were made with AlphaFold2 or AlphaFold-Multimer predicted structures as input. (**A**) Predictions for DB5 target 1JTD. Our method uses one random contact. (**B**) Predictions for RAbD target 2ADF. Our prediction uses four randomly achosen epitope residues. (**C**) Example of high interface and ligand RMSD for an antibody-antigen complex predicted by AlphaFold-Multimer (left). Alignment of predicted chains to the ground truth structure (right). (**D**,**E**) Another example where AlphaFold predicts accurate chain conformations, but incorrect complex. Supplying our method with antigen epitope residues predicted by AlphaFold improves complex prediction quality (left) and CDR loop RMSD (right)

While AlphaFold-Multimer often predicts correct conformations for antibody and antigen chains, the predicted complex can deviate far from the ground truth. For example, Figure S8 (A) shows that although the complex structure is far from the ground truth, the antibody and antigen structures are highly similar to their respective bound counterparts, with less than 2Å complex-RMSD between predicted and unbound antibody chains, and 1.1Å RMSD between predicted and bound antigen chain. In Figure S8 (A,B,D) we provide more examples illustrating this and also show how our model can be used in conjunction with AlphaFold to improve prediction quality when binding site information is known.

### S7 Coordinate Flexibility

Here, we provide further details on maintaining SE(3)-Equivariant updates to rigid frames when some input coordinates are treated as fixed. We note that IPA rigid frames are SE(3)-Equivariant with respect to a single global rigid transformation applied to per-residue local frames [34, Suppl. Material, 1.8.2]. Moreover, the same proof shows that scalar node features are invariant to any global rigid transformation. Thus, setting rigids ***T*** ^(0)^ in the decoder submodule to those derived from a complete set of backbone coordinates results in an SE(3)-Equivariant update to the rigid frames, and an invariant update to the scalar features.

We now argue a more general claim: that IPA can be made SE(3)-Equivariant even when some input coordinates are fixed, flexible or missing. Let *C* = *C*_*fixed*_ ∪ *C*_*flexible*_ ∪ *C*_*missing*_ be a partition of the input residues *i* = 1..*n* denoting those residues with coordinates which should remain fixed, those which are flexible, and those with coordinates that are missing. Without loss of generality, assume that the coordinates which are not missing have mean 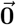, and all missing coordinates are initialized at the origin.

To leave the coordinates corresponding to residues in *C*_*fixed*_ static, we modify the update in Equation (6) to

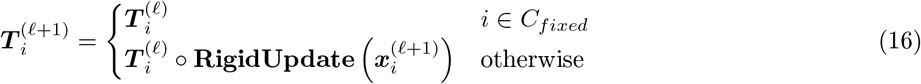

From the equation above, it’s clear that the coordinates are fixed in the output, up to translation. Optionally, we can also replace the prediction of 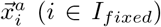 in Equation (7) with the (centered) atom coordinates given as input.

Note that any global rotation applied to the input points will leave the origin fixed, and thus only the fixed or flexible coordinates can change position. The claim of equivariance now follows directly from the equivariance of IPA. To see this, recall that the IPA-layer itself is rotation-equivariant, and that scalar residue features are invariant under the same transformation. Thus, applying a global rotation to all residue coordinates, while keeping the scalar embeddings fixed, will result in only an equivalent update to the local frames.

For practical reasons, mean-centering *all* of the input coordinates does not actually result in an equivariant update – this is because the rigid frames use a specific atom (e.g. *Cα*) to initialize their translation. Thus, in practice, only the rigid translations should have zero-mean.

### S8 CDR-Loop Design

In Section 3.2.1, and Section 4.3, we mention that our architecture is capable of handling direct coordinate information. Moreover, it is possible to treat certain subsets of coordinates as rigid during inference (we actually verify the more general claim - that some coordinates may be fixed, flexible, or missing in Section S7). In settings such as CDR-loop generation, fixing the heavy and light chain framework regions may be practically useful. To enable *de novo* design of loop regions, the CDR L1-L3 and H1-H3 segments can simply be treated as missing. To test whether this approach works in practice, we fine-tuned the same pre-trained model from Section 4.3, while supplying the coordinates of the heavy and light chain framework regions to the structure-decoder module. The framework coordinates are treated as rigid during inference, and the rest of the procedure is implemented exactly as described in Section 4.3. Of course, it is also possible to provide the coordinates of the docked antigen complex in addition to the framework. For example, coordinates on or surrounding the epitope may be treated as flexible, and the others as rigid depending on the use case. We omit this setting here as the manuscript focuses primarily on protein docking.

Fine tuning our model on SAbDab reduces overall sequence perplexity (p = 0.086), and CDR-RMSD (p < 0.005 for CDR H1-H3). We remark that including framework coordinates appears to reduce median CDR H1-H3 RMSD and sequence perplexity, but hypothesis tests comparing our fine-tuned models with and without framework coordinates do not support this claim (p = 0.41, p = 0.43, p = 0.86 for CDR H1, H2, and H3 RMSD). Nevertheless, this outcome provides further empirical justification for our results in Section S7, and acts as a robust proof of concept for how to integrate coordinate information into docking or *de novo* design tasks.

The methods in Table 1 are trained, validated, and tested on different datasets. Because of this, we tried to replicate their training and testing procedures as accurately as possible. To generate our data we use the scheme proposed in Jin et al. [48], generating CDR-clusters at 40% sequence identity and using an 8:1:1 split for training, validation, and test sets respectively. Some example generations are shown in Figure S9

**Table S2:** CDR-loop design with framework coordinates. Results from our method without framework coordinates and without fine tuning (No FT), without framework coordinates and with fine-tuning (FT) and with fine tunng and coordinates for antibody heavy and light chain framework regions (**FT** + **Fr-Coord**). The same criteria and results from our method as described for Section 4.3 are used here.

| Our Method | Structure Prediction |  |  |  | Sequence Prediction |  |  |  |  |
| --- | --- | --- | --- | --- | --- | --- | --- | --- | --- |
|  | RMSD↓ |  |  |  | PPL↓ |  |  | CDR H1-3 |  |
|  | H1 | H2 | H3 | Fr | H1 | H2 | H3 | NSR | PPL |
| No FT | 1.43 | 1.53 | 2.49 | 0.55 | 4.84 | 7.53 | 11.17 | 37.5% | 8.41 |
| FT | 1.11 | 1.04 | 1.88 | 0.82 | 4.46 | 6.71 | 10.68 | 39.7% | 7.67 |
| **FT + Fr-Coord** | 1.03 | 0.98 | 1.78 | – | 4.27 | 6.50 | 10.36 | 40.6% | 7.18 |

**Figure S9:**
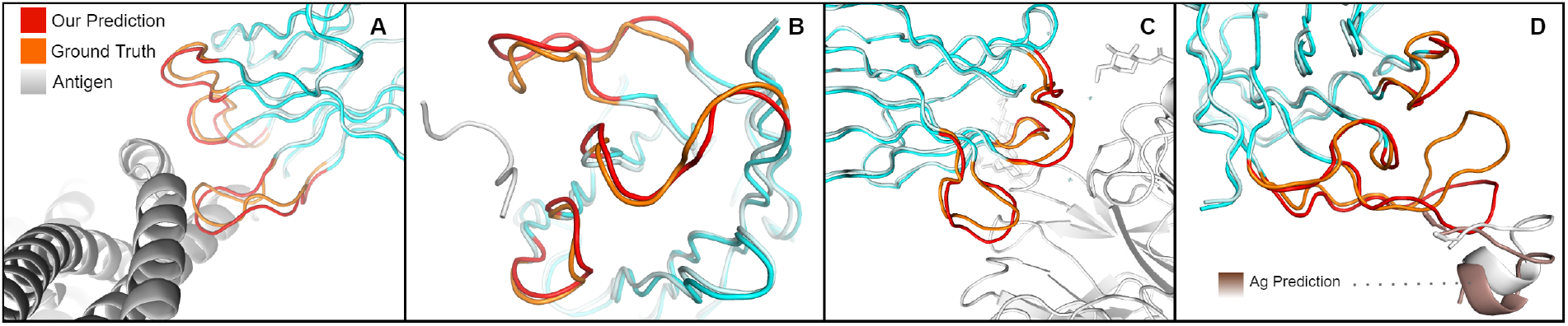
Antibody Docking and CDR Design. Example docking and designs comparing our predictions with native structures. For each example, we give the length (L) of CDR H3 and the RMSD between the predicted (red) and ground truth (orange) conformations. For simplicity, only heavy chains are displayed. Only the bound antigen (gray-white) is shown when the prediction L-RMSD is less than 2 Å. (**A**) Fab of mAb 3E9 in complex with Plasmodium vivax reticulocyte-binding protein 2b (PvRBP2b) (PDB: 6BPA, L = 11, *R*MSD = 1.49). (**B**) Fab of IgG B13I2 bound to synthetic 19-amino acid peptide homolog of the C helix of myohemerythrin (PDB: 2IGF, L = 11, *R*MSD = 1.19). (**C**) Fab of mAb B10 heavy chain in complex with A(H3N2) influenza Virus (PDB: 6N6B, L = 9, *R*MSD = 1.21). (**D**) Fab of igG 7B2 bound to 13-residue HIV-1 GP41 peptide (PDB: 4YDV, L = 17, *R*MSD = 2.86)

### S9 Ablation Studies

We trained several ablated models to identify how different components of our architecture and training procedure contribute to docking performance. We show results for four additional models in Table S3.

We find that removing the shared weight layers and auxiliary FAPE loss from our structure decoder leads to the largest degradation in performance. We also remark that ablating the degree centrality encoding or adding a secondary structure encoding to our input residue features had an insignificant impact on performance. We remark that including ESM1b encodings (+ ESM1b) of each chain did not noticeably improve performance in the blind docking setting. We obtain DockQ scores ≥ 0.23 for 3 targets when ESM1b encodings are used, and 2 targets when the encodings are removed. It appears that these encodings do not significantly improve performance, so we opted for the simpler model instead. Interestingly, the variant of our model which does not use recycling is still able to obtain competitive top-5 performance, but suffers in top-1 performance. Recycling decoder residue features is also competitive with the baseline recycling implementation, but does not result in significantly better performance.

**Table S3:** Ablation Study. We consider the top-1 and top-5 performance of model variants on DB5 unbound targets using 1 contact or 4 interfacial residues as input. This information is randomly sampled independently for each variant, and a total of 15 decoys are generated for each target. Predicted IplDDT is used to rank each decoy. The baseline model is described in the main text. For the four variants we considered removing recycling (No Recycle), adding ESM1b encodings of chain sequences as input (+ESM1b), learning separate weights for each decoder block (No Share Wts), and recycling decoder residue features, rather than encoder residue features (Recycle Dec.). When learning separate weights for decoder layers, we also remove auxiliary FAPE loss.

|  | Top-1 |  |  | Top-5 |  |  | Top-1 |  |  | Top-5 |  |  |
| --- | --- | --- | --- | --- | --- | --- | --- | --- | --- | --- | --- | --- |
|  | IRMS | LRMS | SR | IRMS | LRMS | SR | IRMS | LRMS | SR | IRMS | LRMS | SR |
|  | 4 Interface |  |  |  |  |  | 1-Contact |  |  |  |  |  |
| Baseline | **4.7** | 14.5 | **47.6%** | **3.0** | **8.6** | 69.0% | 5.8 | **13.6** | **45.2%** | 3.5 | 10.2 | <u>59.5%</u> |
| No Recycle | 8.4 | 20.0 | 29.7% | 3.9 | 8.5 | 64.9% | 8.0 | 20.0 | 33.3% | 4.1 | 11.5 | 51.4% |
| + ESM1b | 5.1 | <u>13.8</u> | **47.6%** | <u>3.3</u> | <u>8.8</u> | **73.3%** | **5.6** | <u>18.1</u> | <u>37.7%</u> | <u>3.6</u> | **9.5** | **62.2%** |
| No Share Wts. | <u>4.9</u> | 16.6 | 42.9% | 3.9 | 9.6 | 54.8% | 7.2 | 22.1 | 31.0% | 3.7 | 11.4 | 45.2% |
| Recycle Dec. | 5.3 | **13.7** | 42.9% | 3.4 | 8.8 | **73.3%** | <u>5.7</u> | 18.0 | <u>37.7%</u> | **3.3** | **9.5** | <u>59.5%</u> |

### S10 Data Collection

For all methods, the receptor and ligand chains were randomly rotated and translated before inference. For general proteins, the smaller of the two targets was treated as the ligand (ties broken based on chain order in PDB file). For antibody-antigen chains, the antigen was always treated as the ligand.

Code for EquiDock was downloaded from the author’s github page. Standalone packages for HDock, PatchDock, and ZDock were downloaded from the respective servers. For HDock and PatchDock, all binding interface and contact information was given as input. Still, results required an additional post-processing step when run locally. For this, we enumerate all predictions of each program and choose the lowest energy prediction satisfying the interface and contact criteria. We reiterate that interface and contacts are defined using *Cα* atoms with 10Å cutoff. In some cases, HDock or PatchDock did not produce any decoys meeting all criteria. In these cases, we choose the lowest scoring model with the most recovered interface residues and contacts.

AlphaFold and AlphaFold-Multimer were run with ColabFold [90] using the provided template and MSA servers. Default settings were used for all other options. ColabFold’s monomer setting was used to predict all chains in the DB5 benchmark, and all antigen chains in the RAbD and Ab-Bench benchmarks. The multimer setting was used to generate all predicted antibody structures with bound heavy and light chains.

As mentioned in Section 3.2.2, we filter unbound and predicted targets based on RMSD to the bound conformation. Full lists of targets used for comparisons is included with the code at https://github.com/MattMcPartlon/proteindocking

### S11 Extended Results and Examples

#### S11.1 Docking Benchmark Version 5

**Table S4:**
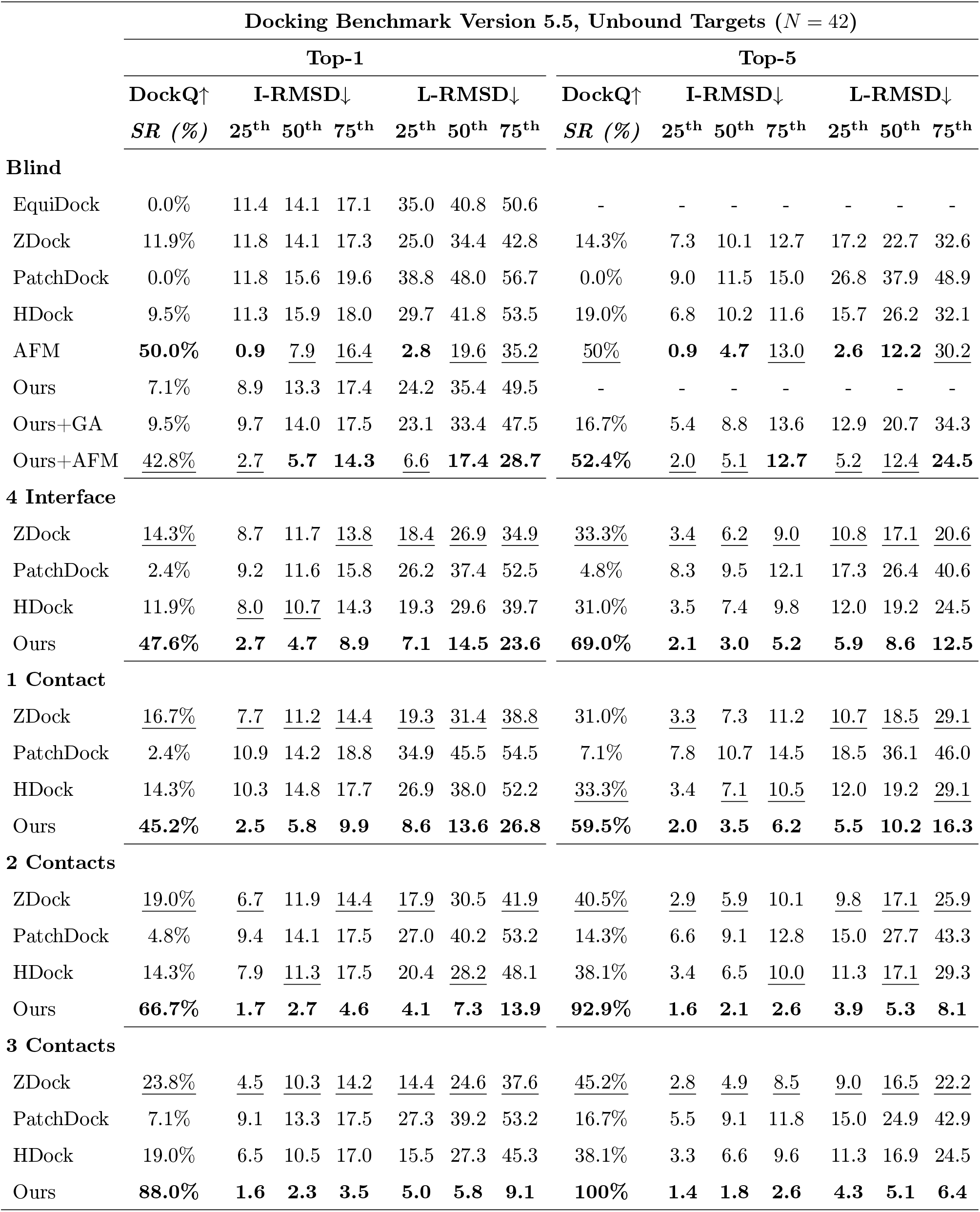
Results for DB5 Unbound Targets.

**Table S5:** Results for DB5 Predicted Targets.

| Docking Benchmark Version 5.5, Predicted Targets ($N = 22$) | | | | | | | | | | | | | | |
| --- | --- | --- | --- | --- | --- | --- | --- | --- | --- | --- | --- | --- | --- | --- |
|  | Top-1 |  |  |  |  |  |  | Top-5 |  |  |  |  |  |  |
| | DockQ↑ | I-RMSD↓ | | | L-RMSD↓ | | | DockQ↑ | I-RMSD↓ | | | L-RMSD↓ | | |
|  | *SR (%)* | 25<sup>th</sup> | 50<sup>th</sup> | 75<sup>th</sup> | 25<sup>th</sup> | 50<sup>th</sup> | 75<sup>th</sup> | *SR (%)* | 25<sup>th</sup> | 50<sup>th</sup> | 75<sup>th</sup> | 25<sup>th</sup> | 50<sup>th</sup> | 75<sup>th</sup> |
| **Blind** |  |  |  |  |  |  |  |  |  |  |  |  |  |  |
| EquiDock | 0.0% | 9.8 | **12.2** | **15.4** | 26.4 | 42.1 | <u>50.1</u> | - | - | - | - | - | - | - |
| ZDock | **9.1%** | **8.5** | <u>12.9</u> | <u>17.6</u> | **19.5** | **29.5** | **34.8** | **18.2%** | **5.5** | **10.0** | 13.0 | **12.0** | **22.4** | **31.1** |
| PatchDock | 4.5% | 11.5 | 14.4 | 19.4 | 29.8 | 38.2 | 55.5 | 9.1% | <u>8.4</u> | <u>10.2</u> | <u>11.6</u> | 18.6 | <u>27.4</u> | <u>32.0</u> |
| HDock | **9.1%** | 12.0 | 15.2 | 20.6 | 27.5 | 36.9 | 63.6 | **18.2%** | 8.6 | 11.0 | 13.3 | <u>17.0</u> | 28.6 | 34.1 |
| Ours | **9.1%** | <u>9.6</u> | 13.8 | 18.9 | <u>22.8</u> | <u>34.3</u> | 58.3 | - | - | - | - | - | - | - |
| **4 Interface** |  |  |  |  |  |  |  |  |  |  |  |  |  |  |
| ZDock | 9.1% | <u>7.2</u> | 11.9 | 15.3 | <u>17.6</u> | <u>27.8</u> | <u>33.1</u> | <u>31.8%</u> | <u>2.9</u> | <u>6.8</u> | 10.2 | <u>8.1</u> | <u>15.2</u> | 26.9 |
| PatchDock | 4.5% | 9.7 | 11.7 | <u>14.4</u> | 21.1 | 29.5 | 38.6 | 18.2% | 5.1 | 8.3 | <u>10.0</u> | 15.1 | 18.1 | 27.8 |
| HDock | 9.1% | 8.7 | <u>11.3</u> | <u>14.4</u> | 21.1 | 30.1 | 41.4 | 27.3% | 3.6 | 8.3 | 10.1 | 14.6 | 19.6 | <u>25.9</u> |
| Ours | **59.1%** | **2.2** | **3.2** | **7.4** | **6.1** | **7.1** | **22.2** | **68.2%** | **2.2** | **2.6** | **4.1** | **5.4** | **6.8** | **10.6** |
| **1 Contact** |  |  |  |  |  |  |  |  |  |  |  |  |  |  |
| ZDock | <u>13.6%</u> | <u>6.5</u> | <u>12.0</u> | <u>17.6</u> | <u>17.0</u> | <u>28.4</u> | <u>37.8</u> | <u>36.4%</u> | <u>2.7</u> | <u>6.5</u> | 13.0 | <u>8.0</u> | <u>18.4</u> | <u>31.1</u> |
| PatchDock | 4.5% | 10.0 | 14.4 | 19.1 | 21.1 | 34.6 | 54.9 | 27.3% | 4.0 | 8.4 | <u>11.2</u> | 13.1 | 19.0 | 32.0 |
| HDock | 9.1% | 10.9 | 15.0 | 20.6 | 27.5 | 39.0 | 63.6 | 22.7% | 6.0 | 10.2 | 12.8 | 16.8 | 23.8 | 33.5 |
| Ours | **27.3%** | **3.6** | **7.1** | **10.3** | **10.0** | **18.3** | **29.1** | **54.5%** | **2.4** | **2.8** | **7.2** | **6.7** | **10.5** | **15.7** |
| **2 Contacts** |  |  |  |  |  |  |  |  |  |  |  |  |  |  |
| ZDock | 9.1% | <u>7.2</u> | <u>11.5</u> | 15.6 | <u>17.0</u> | <u>26.9</u> | <u>34.8</u> | <u>36.4%</u> | <u>2.4</u> | <u>6.5</u> | 13.0 | <u>5.8</u> | <u>17.3</u> | 31.1 |
| PatchDock | 4.5% | 9.1 | 12.8 | <u>15.4</u> | 22.1 | 31.1 | 50.1 | 27.3% | 4.0 | 7.9 | <u>10.6</u> | 13.1 | 18.6 | 32.0 |
| HDock | <u>13.6%</u> | 7.9 | 14.6 | 20.6 | 23.1 | 39.0 | 63.6 | 22.7% | 6.0 | 8.8 | 11.1 | 16.8 | 24.0 | <u>29.5</u> |
| Ours | **66.7%** | **2.3** | **3.3** | **6.4** | **6.4** | **7.3** | **17.7** | **90.5%** | **1.9** | **2.4** | **3.0** | **5.1** | **6.5** | **7.3** |
| **3 Contacts** |  |  |  |  |  |  |  |  |  |  |  |  |  |  |
| ZDock | 13.6% | <u>6.5</u> | <u>11.0</u> | <u>15.6</u> | <u>15.2</u> | <u>28.4</u> | <u>34.8</u> | <u>40.9%</u> | <u>2.4</u> | <u>6.9</u> | 13.2 | <u>5.8</u> | <u>15.2</u> | 31.4 |
| PatchDock | 4.5% | 9.8 | 13.7 | <u>15.6</u> | 27.1 | 31.4 | 52.9 | 27.3% | 3.7 | 7.3 | <u>9.9</u> | 11.1 | 16.7 | 32.0 |
| HDock | <u>18.2%</u> | 7.8 | 15.0 | 20.6 | 18.7 | 38.4 | 63.6 | 27.3% | 3.8 | 8.4 | 10.1 | 14.6 | 22.7 | <u>28.9</u> |
| Ours | **75.0%** | **1.5** | **2.4** | **4.1** | **4.6** | **7.8** | **11.7** | **95.0%** | **1.4** | **2.0** | **2.4** | **4.0** | **5.3** | **8.0** |

#### S11.2 Antibody Benchmark

**Table S6:**
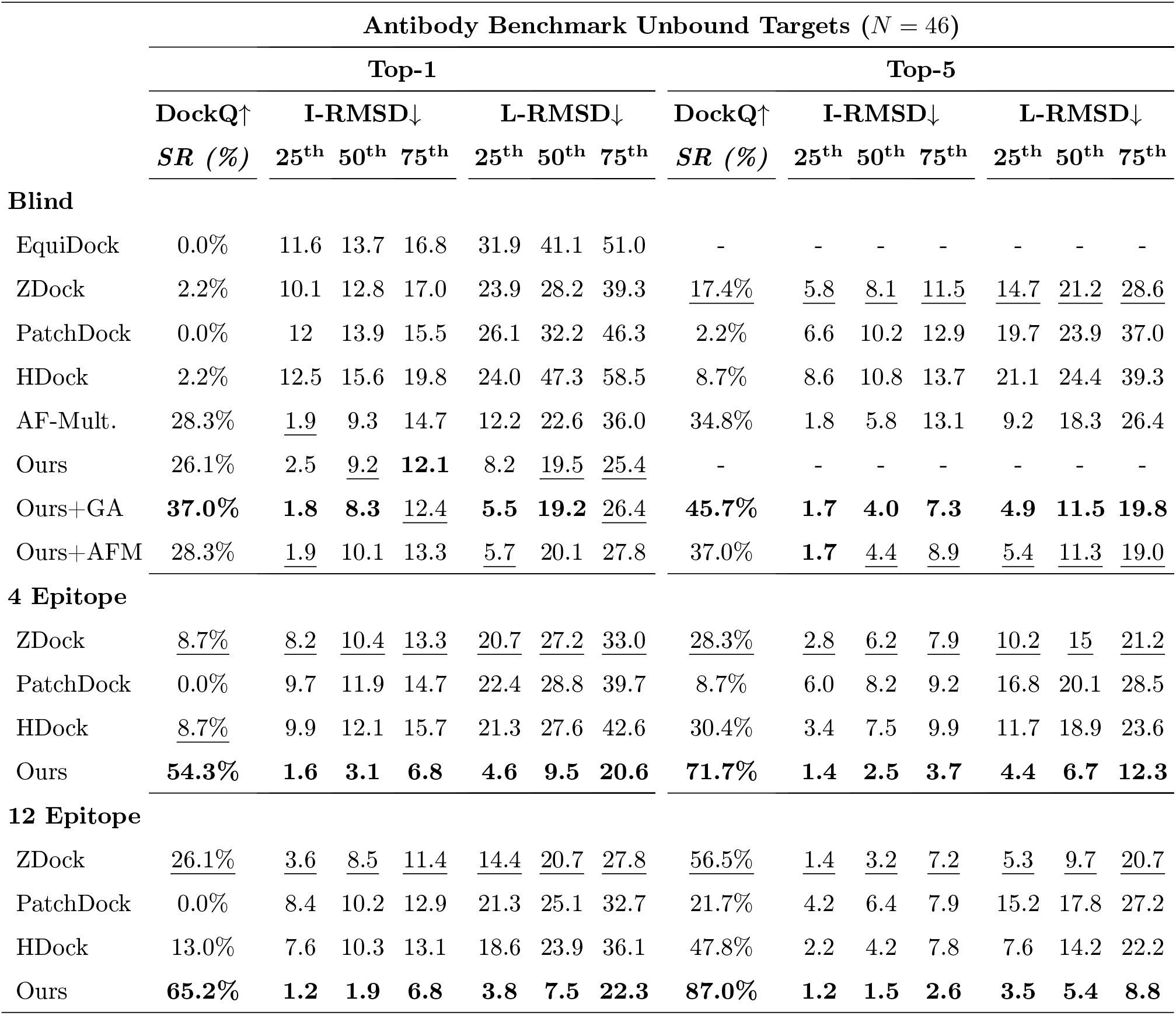
Results for Antibody Benchmark Unbound Targets.

**Table S7:** Results for Antibody Benchmark Predicted Targets.

| Antibody Benchmark Predicted Targets ($N = 26$) | | | | | | | | | | | | | | |
| --- | --- | --- | --- | --- | --- | --- | --- | --- | --- | --- | --- | --- | --- | --- |
|  | Top-1 |  |  |  |  |  |  | Top-5 |  |  |  |  |  |  |
| | DockQ↑ | I-RMSD↓ | | | L-RMSD↓ | | | DockQ↑ | I-RMSD↓ | | | L-RMSD↓ | | |
|  | *SR (%)* | 25<sup>th</sup> | 50<sup>th</sup> | 75<sup>th</sup> | 25<sup>th</sup> | 50<sup>th</sup> | 75<sup>th</sup> | *SR (%)* | 25<sup>th</sup> | 50<sup>th</sup> | 75<sup>th</sup> | 25<sup>th</sup> | 50<sup>th</sup> | 75<sup>th</sup> |
| **Blind** |  |  |  |  |  |  |  |  |  |  |  |  |  |  |
| EquiDock | 0.0% | 13.2 | 14.5 | 16.4 | 38.3 | 41.6 | 50.0 | - | - | - | - | - | - | - |
| ZDock | 3.8% | 10.6 | 12.9 | 15.4 | 21.9 | 26.8 | 37.3 | <u>7.7%</u> | <u>6.4</u> | <u>9.6</u> | <u>11.5</u> | <u>14.7</u> | <u>19.8</u> | 28.7 |
| PatchDock | 0.0% | 12.3 | 14.0 | 18.8 | 26.3 | 33.3 | 49.5 | 3.8% | 7.8 | 10.9 | 12.2 | 18.4 | 22.7 | <u>28.0</u> |
| HDock | 0.0% | 11.9 | 13.5 | 18.9 | 25.5 | 31.2 | 52.4 | 3.8% | 8.1 | 10.1 | 12.4 | 18.3 | 25.2 | 30.7 |
| Ours | <u>26.9%</u> | <u>2.8</u> | <u>10.4</u> | <u>13.9</u> | <u>9.4</u> | <u>22.9</u> | <u>26.4</u> | - | - | - | - | - | - | - |
| Ours+GA | **42.3%** | **1.9** | **7.3** | **9.0** | **6.5** | **15.5** | **19.4** | **46.2%** | **1.8** | **7.2** | **9.1** | **6.5** | **13.5** | **17.3** |
| **4 Epitope** |  |  |  |  |  |  |  |  |  |  |  |  |  |  |
| ZDock | <u>7.7%</u> | <u>7.4</u> | <u>9.4</u> | <u>13.2</u> | <u>18.6</u> | <u>24.7</u> | <u>31.8</u> | <u>30.8%</u> | <u>3.9</u> | <u>6.1</u> | <u>8.8</u> | <u>8.9</u> | <u>15.3</u> | <u>21.0</u> |
| PatchDock | 3.8% | 10.3 | 12.2 | 14.5 | 24.8 | 27.2 | 40.0 | 7.7% | 6.0 | 7.7 | 9.5 | 15.2 | 19.4 | 26.4 |
| HDock | 3.8% | 9.0 | 13.2 | 14.1 | 23.0 | 27.6 | 35.2 | 19.2% | 5.0 | 7.5 | 10.4 | 15.8 | 19.5 | 25.9 |
| Ours | **53.8%** | **1.9** | **2.7** | **9.3** | **4.8** | **8.5** | **28.0** | **69.2%** | **1.5** | **2.4** | **3.4** | **4.4** | **6.0** | **11.8** |
| **12 Epitope** |  |  |  |  |  |  |  |  |  |  |  |  |  |  |
| ZDock | <u>19.2%</u> | <u>5.7</u> | 10.3 | <u>11.6</u> | <u>14.0</u> | <u>23.2</u> | <u>28.6</u> | <u>57.7%</u> | <u>2.0</u> | <u>3.9</u> | <u>6.2</u> | <u>4.8</u> | <u>11.8</u> | <u>15.8</u> |
| PatchDock | 11.5% | 7.4 | <u>10.2</u> | 12.4 | 17.3 | 26.1 | 38.9 | 30.8% | 4.5 | 6.0 | 7.4 | 8.9 | 17.2 | 24.3 |
| HDock | 7.7% | 7.8 | <u>10.2</u> | 13.5 | 17.9 | 24.3 | 30.2 | 50.0% | 2.1 | 5.3 | 8.6 | 5.8 | 13.2 | 22.7 |
| Ours | **69.2%** | **1.3** | **1.8** | **5.8** | **3.7** | **6.1** | **21.1** | **88.5%** | **1.3** | **1.7** | **3.2** | **3.7** | **4.8** | **9.8** |

#### S11.3 Rosetta Antibody Design

**Figure S10:**
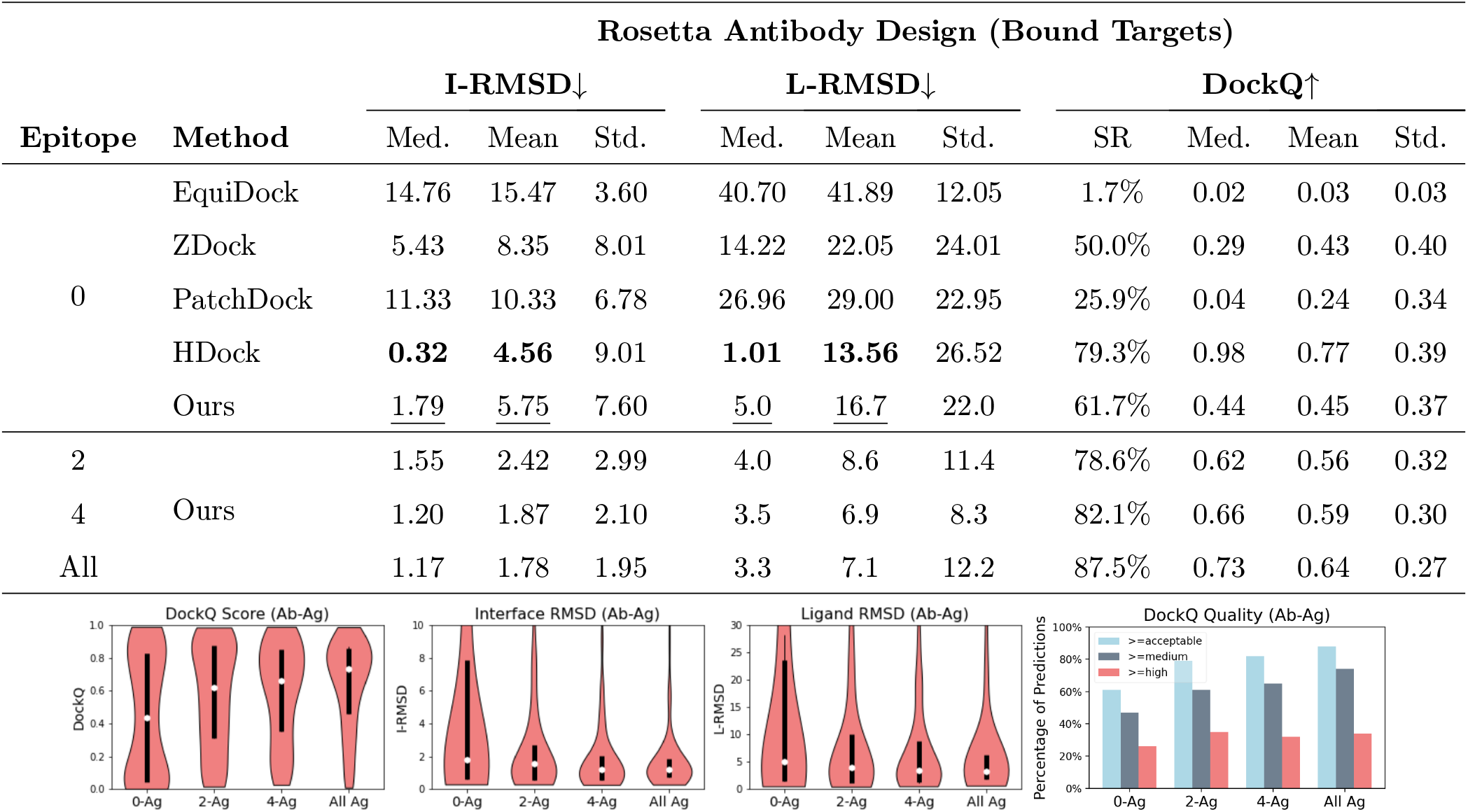
Rosetta Antibody Design Bound Targets. Results on the RAbD test set using bound chains as input to each docking method. Results for our method are generated after fine-tuning on bound antibody-antigen chains. The *x*-axis in the below 4 pictures show the number of epitope residues provided to the docking methods. DockQ score cutoffs for acceptable, medium and high quality predictions are ≥ 0.23, ≥ 0.49, and ≥ 0.80

**Figure S11:**
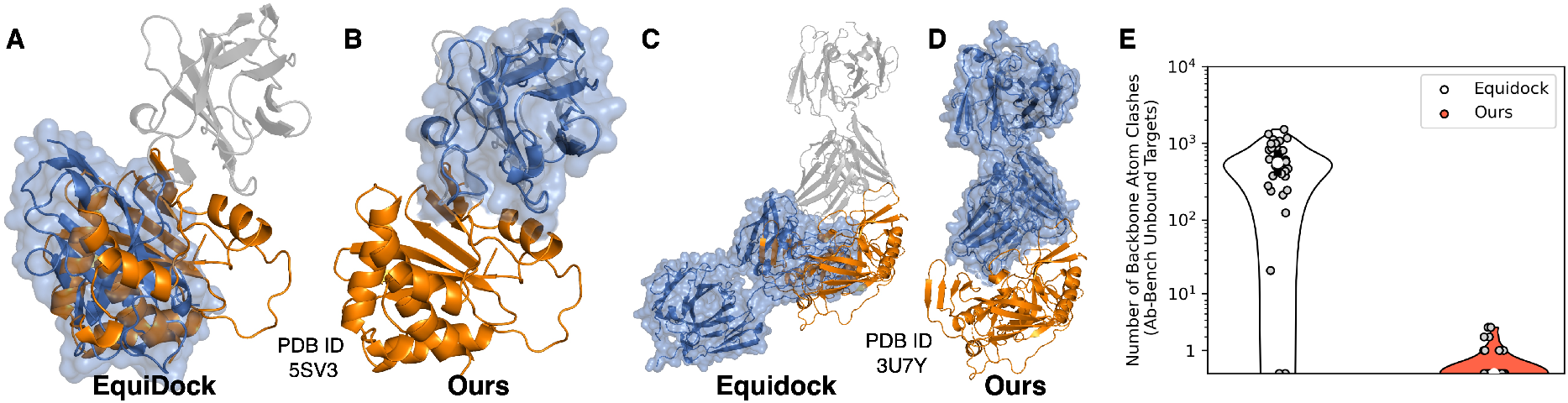
Comparison Between Our Method and Equidock. Blind docking predictions for a single domain antibody targeting the toxin Ricin (**A** and **B**), and therapeutic antibody which targets the CD4 binding site on the HIV-1 spike protein (**C** and **D**). In (A–D), we show the bound antigen in orange and bound antibody in light gray. For clarity we align each complex prediction to the ground truth using only the antigen chain, and show only the predicted antibody in blue. We also show the solvent accessible surface of antibody predictions (independent of the antigen) to better illustrate surface intersections. For both of these targets, the RMSD between bound and unbound antigen chains is less than 2Å. (**E**) Distribution of the number of steric clashes for blind docking DB5 unbound targets. We consider only backbone atom clashes, since EquiDock cannot modify side-chain conformations. Two atoms are said to clash if each atom belongs to a different chain, and the pairwise distance is less than 90% the sum of their van der Waals radii.

**Figure S12:**
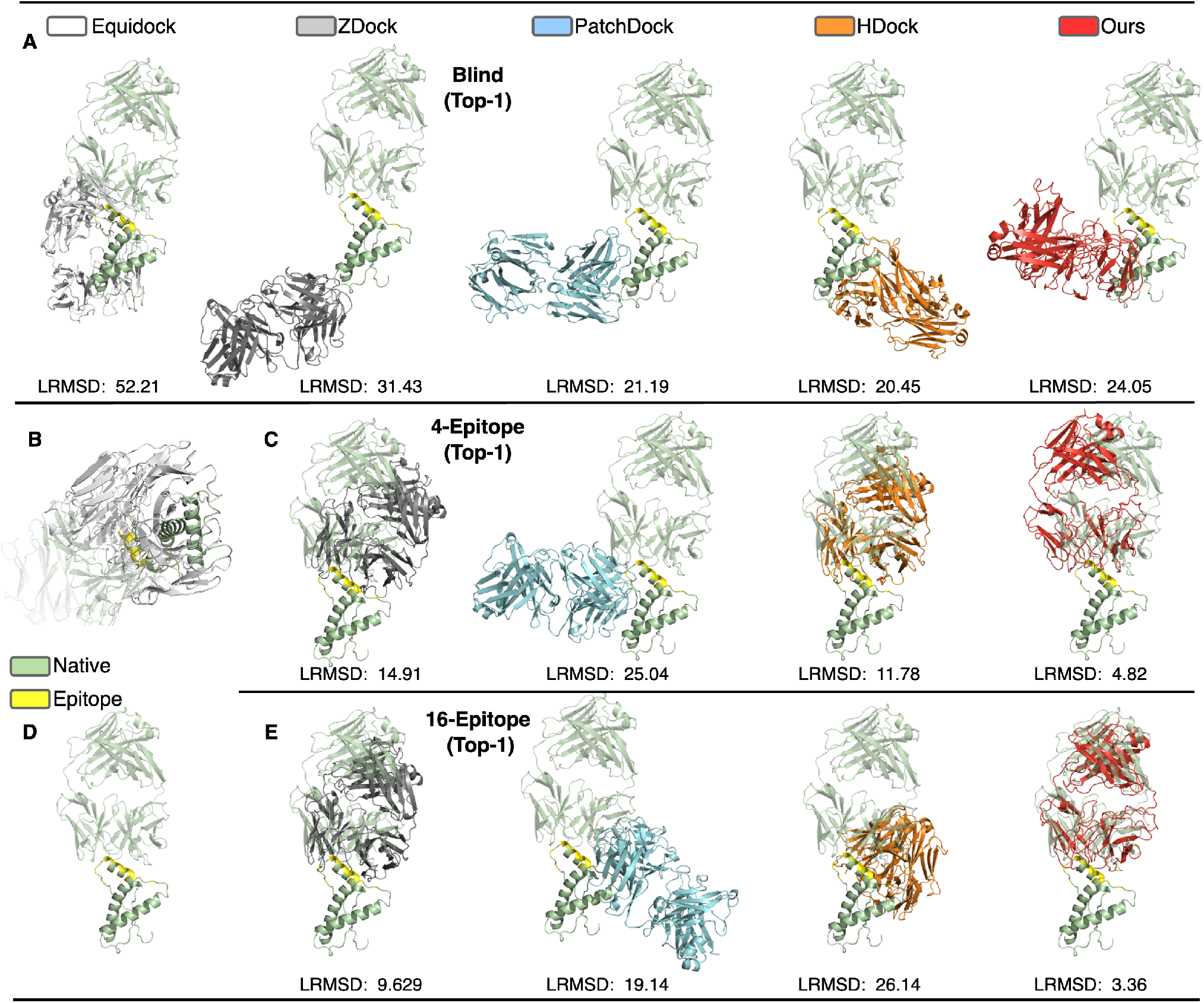
Docking Predictions for Antibody Benchmark Target 2W9E. Protein backbones are shown in cartoon with ground-truth antibody and antigen structures shown in green for each figure. The antigen epitope is highlighted in yellow. We show the predicted antibody orientation relative to the ground truth antigen in a separate color for each method. Ligand RMSD (LRMSD) is shown for each prediction. (**A**) Blind docking predictions for methods EquiDock, ZDock, PatchDock, HDock, and DockGPT. (**B**) Close up of EquiDock’s prediction showing excessive surface overlap between antibody and antigen chain predictions. (**C**) Top-1 docking predictions for each method, except EquiDock given four epitope residues. (**D**) Ground truth complex. (**E**) Top-1 docking predictions for each method, except EquiDock given 12 epitope residues.

## Footnotes

1 Downloaded from https://github.com/drorlab/DIPS.

2 See [here](http://www.bioinf.org.uk/abs/info.html) for a nice summary of CDR numbering schemes and changes in corresponding CDR loop definitions over time.

3 DockQ is publicly available for download at http://github.com/bjornwallner/DockQ/

